# The targeting of non-fibrillar polyQ via distinct VCP-proteasome coupling

**DOI:** 10.1101/2025.04.01.646186

**Authors:** Dorothy Yanling Zhao, Chenchen Mi, Jonathan Wagner, Wenghong Jiang, Itika Saha, Matthias Poege, Peng Xu, Juergen Plitzko, Qiang Guo, Florian Beck, Wolfgang Baumeister

## Abstract

The aggregation of proteins containing expanded poly-glutamine (polyQ) repeat sequences is a cytopathological hallmark of several dominantly inherited neurodegenerative diseases (ND), including Huntington’s disease (HD). Previously, we observed that long polyQ repeats form a fibrillar core, surrounded by amorphous and soluble intermediates, which compared to the fibrils, are more readily taken up by autophagosomes ^1^. In this study, we investigated how the alternative major degradation pathway in cultured cells, ubiquitin proteasome system (UPS), interacts with the different pools of polyQ.

We observed that the AAA+ ATPase, valosin-containing protein VCP/p97, in collaboration with proteasomes, plays a crucial role in degrading non-fibrillar polyQ. As both VCP and proteasomes were recruited to the polyQ intermediates peripheral to the fibrils, we imaged these regions by *in situ* cryo-ET for subtomogram averaging. VCP predominantly adopts an ATP-bound state, often in an active processing conformation with resolved NPLOC4-like cofactor density. The region was also enriched with the 26S proteasome (26S), 20S proteasome core (20S), and 19S regulatory particles (19S). Distance analysis of the macromolecules revealed a striking proximity between VCP and the 20S *in situ.* This is likely mediated by VCP’s C-terminal hydrophobic-tyrosine-X (HbYX) motif, which binds the 20S via hydrophobic interactions, as confirmed by cryo-EM single-particle analysis (SPA) and *in vitro* assays. Our study highlights a distinct VCP-20S coupling mechanism, where VCP functionally overlaps with the 19S complex in substrate unfolding, facilitating the degradation of polyQ intermediates by the 20S catalytic core.

## Introduction

Numerous neurodegenerative disorders (NDs), including Alzheimer’s disease (AD), Parkinson’s disease, frontotemporal dementia (FTD), amyotrophic lateral sclerosis (ALS), and Huntington’s disease (HD), are associated with the formation of toxic aggregates resulting from protein misfolding ^2-5^. HD, characterized by progressive motor and cognitive decline, is the most common member of a group of NDs caused by dominantly inherited polyglutamine (polyQ) repeat expansions in otherwise unrelated proteins ^6-8^. Expansion of the polyQ sequence in exon 1 of the huntingtin (Htt) protein beyond approximately 37Q is a strong predictor of HD ^6,7,9,10^. The length of the polyQ tract correlates positively with aggregation propensity and disease severity, while inversely correlating with the age of disease onset ^6,7^. Longer repeats (up to >100 Q) more readily form amyloid-like fibrils, leading to the formation of highly stable inclusion bodies ^11,12^, which initially appear in striatal neurons and later spread to other brain regions as the pathology progresses ^6,7,11^.

PolyQ-expanded Htt aggregates exhibit structurally distinct forms, including dynamic soluble oligomers and stable fibrils with a cross-β structure ^12-19^. The amorphous polyQ has been identified as an intermediate phase for fibril formation ^20^, shown to accumulate at the periphery of the fibrillar core and solidify upon autophagy inhibition ^1^. While the large fibrils are considered less toxic ^21-23^, the soluble oligomers are recognized as major toxic agents due to their ability to aberrantly interact with cellular machineries ^14,18,24-27^. Together, both species contribute to cytopathology by sequestering key cellular proteins and physically disrupting subcellular membrane structures ^1,13-15,26,28^.

Mammalian cells, including neurons, utilize autophagy and the ubiquitin-proteasome system (UPS) to clear misfolded and aggregated substrates, thereby maintaining of cellular proteostasis ^29-33^. Both pathways depend on substrate poly-ubiquitinylation as a signal for degradation. Our cryo-ET study has shown that autophagy has a limited capacity to clear polyQ aggregates ^1^. The alternative UPS involving the proteasome, is also dysregulated in a wide range of proteinopathies including HD ^33-39^, as it faces major challenges in degrading stable aggregates due to steric hindrance ^32,40-45^. The 26S proteasome, responsible for ubiquitin-mediated protein degradation, is composed of the 20S catalytic core and the 19S regulatory particle. The 20S consists of seven α and seven β subunits arranged in a conserved cylindrical structure with an α7β7β7α7 stacking. The 19S plays a critical role in recognizing and unfolding polyubiquitinated substrates, opening the 20S core, and translocating unfolded proteins into the 20S catalytic chamber ^46-50^.

Chaperone-mediated substrate unfolding often serves as a prerequisite upstream of proteasome degradation ^51-54^. The 540 kDa AAA+ ATPase VCP, together with its ubiquitin-selective cofactor complex NPLOC4/UFD1, can facilitate the disassembly of the microtubule-associated Tau protein aggregates ^51,55^. Over 30 human VCP mutations have been identified and are associated with dominantly inherited neurodegenerative diseases, including vacuolar tauopathy, inclusion body myopathy, ALS, and FTD ^55-59^. While loss-of-function mutations disrupt proteostasis, hyperactive VCP mutations ^59-62^ can also drive disease, potentially by generating misfolded intermediates that promote neurotoxic spreading ^51^. A deeper understanding of VCP’s function *in situ* remains critical to the field of proteostasis.

Here we address the roles of VCP and proteasome in the context of HD aggregates using *in situ* cryo-ET and subtomogram averaging ^63^. Most notably, we observe that VCP is proximal to not only to the 26S but also to the 20S *in situ*. Confirmed by *in vitro* assays and cryo-EM SPA, we reveal a distinct VCP-20S coupling mechanism in higher eukaryotic cells, capable of degrading non-fibrillar polyQ intermediates.

## Results

### 1. Non-fibrillar polyQ intermediates are cleared by VCP and proteasome

To investigate the roles of VCP and the proteasome in polyQ clearance in neuronal cells, we expressed ecdysone-regulated Htt exon 1 GFP fusion proteins with 64Q repeats ^14,64^. After 48 h of induction with muristerone A, the majority of 64Q appeared diffused in the cell, with approximately 10% of cells exhibited small foci of varying brightness ^1,14^. We assessed the impact of VCP and proteasome inhibition on the clearance of these two states by applying pharmacological inhibitors NMS873 and MG132 for 12 h after inducer removal. Total polyQ levels were assessed by immunoblotting, while SDS-resistant aggregated fibrils were analyzed by slot blotting. The analysis revealed an approximate two-fold increase in the total 64Q pool following single or combinatorial inhibition, though SDS-resistant aggregate levels did not consistently increase (Figure 1A). These findings suggest that VCP and the proteasome are involved in the degradation of non-fibrillar, SDS-soluble polyQ.

**Figure 1.**
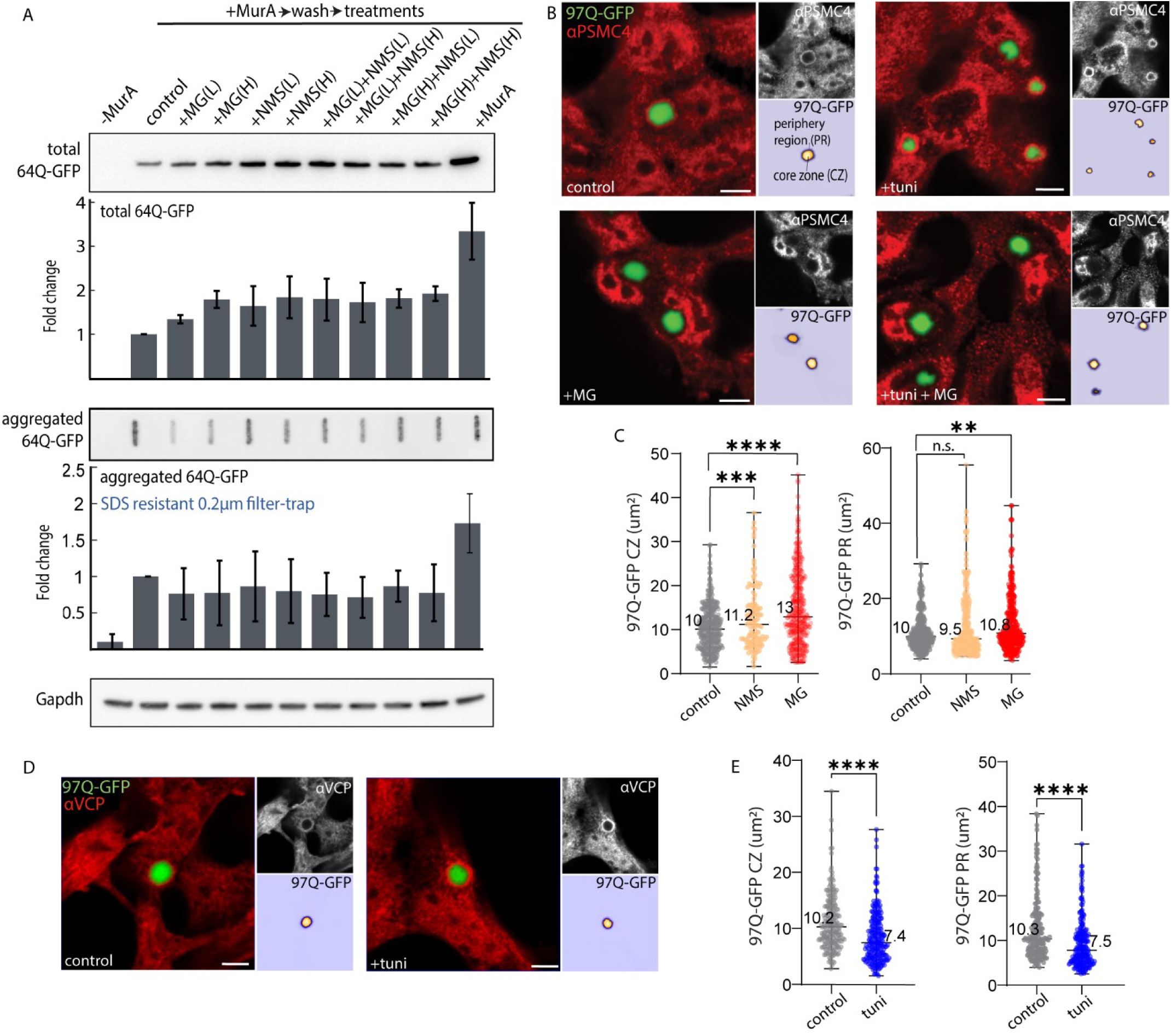
Non-fibrillar polyQ intermediates are targeted by VCP and proteasome. **(A**) Immunoblot and slot blot analyses with the indicated antibodies of total and aggregated 64Q-GFP from Neuro2a cells following 2-day muristerone A induction and removal (control) or subsequent 12 h treatments to inhibit VCP (NMS873: L= 5 μM, H = 10 μM) and/or proteasome (MG132: L = 5 μM, H = 10 μM). Gapdh used as a loading control. Also included are -MurA: no induction, and +MurA: muristerone A induction without removal. Quantification normalized to muristerone A removal without further treatments (control = 1), error bars: s.d., n > 3. **(B, D)** Representative confocal images of 97Q-GFP control or treated HEK293 cells, co-stained with PSMC4 for proteasome (**B**) or VCP (**D**). Also included are 9 h treatments with MG132 (10 μM) to inhibit proteasome and/or tunicamycin (2 μg/ml) to induce ER stress. (**C**) Image quantification of 97Q-GFP in the central zone (CZ) and peripheral region (PR) under control or following 9 h treatment of NMS873 (10 μM) or MG132 (10 μM) (CZ: control: n = 332, NMS873: n = 119, MG132: n = 393; PR: control: n = 413, NMS873: n = 373, MG132: n = 357). **(E**) Image quantification of 97Q-GFP in the CZ and PR under control or following 9 h tuni treatment (CZ: control: n = 206, tuni: n = 232; PR: control: n = 179, tuni: n = 229) Median values are displayed (**C, E**), intensity standardized with threshold 800-8000 for PR, and > 8000 for CZ, **** p < 0.0001, *** p < 0.001, ** p < 0.01, p values generated from two-tailed student’s t-test. Scale bars: 5 μm (**B**, **D**).

To assess VCP and proteasome enrichment across different polyQ phases, we isolated 150Q, which is expressed similarly to 64Q in Neuro2a cells but is highly enriched in the aggregated form ^1^. We analyzed total 150Q and filter-partitioned aggregated 150Q (>0.2 μm) from lysates, enriched via GFP pull-down (GFP-trap) for label-free quantitative mass spectrometry (Figure S1A-E). Aggregated 150Q showed enrichment of 19S subunits and associated receptors such as UBQLNs, which mediate ubiquitin recognition ^34^ (Figure S1A, S1C). VCP and its NPLOC4/UFD1 cofactor complex, known to interact with disease aggregates including polyQ ^25,51,65^ were also enriched (Figure S1A). Notably, 20S subunits was not enriched with aggregated 150Q. In contrast, total 150Q was enriched with VCP, and both 19S and 20S subunits, including PSMA1/A5/A6/A7 (Figure S1B-C). These findings suggest differential engagement of 20S and 19S with distinct polyQ species. The dissociation of 26S into 20S and 19S, potentially regulated by Ecm29 detected in our system (Figure S1A-B), may occur readily under a range of cellular stress ^39,66-68^. The selective enrichment of 19S subunits in aggregated 150Q likely reflect binding and sequestration rather than active degradation ^40^. Together, these data indicate that while VCP and the 19S proteasome subcomplex associate with the aggregated polyQ, their degradative effect primarily targets the SDS-soluble polyQ intermediates.

### 2. VCP and proteasome co-localize with polyQ intermediates at fibrillar periphery

The Htt polyQ protein undergoes a liquid-to-solid phase transition, driving aggregation towards SDS-resistant amyloid-like fibrils, accompanied by an increase in polyQ-GFP fluorescence ^1,20^. To better visualize VCP and proteasome with respect to the polyQ aggregates, we used confocal fluorescence microscopy with 97Q-GFP (97Q) in HEK293 cells, which exhibit lower background staining and enable reliable quantifications ^1^. Fluorescence microscopy shows that 97Q and 150Q aggregates are similar in brightness and prevalence ^1^, both contain a highly fluorescent central zone (CZ) in equilibrium with a dim peripheral region (PR), where amorphous and soluble polyQ intermediates accumulate (Figure 1B). Quantification of cross-sectional areas at the aggregate equator, with PR intensity thresholded to approximately one-tenth of CZ, revealed that both zones expanded after 9 h of proteasome (MG132) and VCP (NMS873) inhibition, with proteasome inhibition having a stronger effect (Figure 1C). Consistent with their role in degrading non-fibrillar polyQ intermediates, proteasome and VCP concentrated in the PR zone (Figure 1B, 1D), with proteasome enrichment lost upon its inhibition by MG132 (Figure 1B).

Given the observed ER enrichment around polyQ aggregates ^13,25,65^ (Figure S1F), we hypothesized that enhancing ER stress to activate ER-associated degradation may recruit additional proteasomes and VCP to the ER-proximal polyQ aggregates, thereby counteracting their growth. To test this, we induced ER stress ^25^ with tunicamycin (tuni) for 9 h, which reduced aggregate size in both the CZ and PR zones (Figure 1E) and correlated with enhanced VCP and proteasome recruitment (Figure 1B, 1D). Quantification of VCP (Figure S1G) and proteasome as indicated by PSMB5 staining for the 20S and PSMC4 staining for the 19S (Figure S1H-I) within a 1.5 μm radius around the CZ, confirmed increased recruitment upon ER stress. To demonstrate that the tunicamycin effect occurs via the UPS, we repeated the measurement in Hek293 LC3 A/B knockout cells, in which autophagy is abolished ^1^, and observed a similar trend (Figure S1J). Our drug treatments, including their concentration and duration, did not adversely affect cell viability, as assessed by live Annexin V staining (Figure S1K). Altogether, these data (Figure 1) demonstrate that VCP and proteasomes degrade polyQ intermediates and localize to the aggregate periphery, where these intermediates accumulate. Further induced ER stress can enhance VCP and proteasome recruitment to ER-proximal aggregates.

### 3. *In situ* visualization of VCP and proteasomes around polyQ aggregates

To elucidate the macromolecular interactions around the polyQ aggregates, we employed cryo-correlated light and electron microscopy (cryo-CLEM) ^69-71^ coupled with subtomogram averaging ^72^ (Figure S2A). PolyQ-expressing cells were cultured, treated on cryo-EM grids, and vitrified by plunge freezing. Tunicamycin was included (9 h) to enhance VCP and proteasome densities around the aggregates, reducing the number of tomograms needed to achieve reasonable resolution in the reconstructions. GFP signal, identified through cryo-CLEM, guided aggregate targeting during lamella preparation with focused ion beam (FIB) milling. Inspection of the lamella on a 300kV TEM (Figure S2D, S3C, S4C) revealed the aggregates, enabling tilt-series collection focused on the peripheral region of the aggregates.

We conducted initial analysis using a dataset acquired with a pixel size of 3.35 Å, binned by a factor of two (Figure 2 and Figure S2-5). To reduce false positives in template matching and subtomogram extraction, regions above and below the lamella, tomogram edges, ice crystals, organelles and membranes were removed. The removal of membrane regions also prevents the selection of membrane-bound macromolecules for downstream analysis. The coordinates were aligned and classified using Stopgap ^72^ to eliminate incorrect coordinates that did not converge to accurate class averages. After iterative classifications and alignments, the coordinates of the macromolecules were mapped back to the tomographic volume for visual inspection (Figure 2A-G, S3D-J, S4D-J). This approach enabled a resolution of 13.7 Å for ribosomes as a benchmark (Figure S5A-B). Initial template matching of the 26S was performed using a low-pass filtered truncated reference to recover full structure from the dataset. A similar approach did not recover double-capped proteasomes (30S), suggesting low abundance. As many barrel-like structures could not be assigned to either 26S or 30S, they were instead template-matched to the 20S. For 20S, 19S, and different states of VCP, known PDB structures were low-pass filtered to 40 Å as templates for the initial matching (Figure S5A).

**Figure 2.**
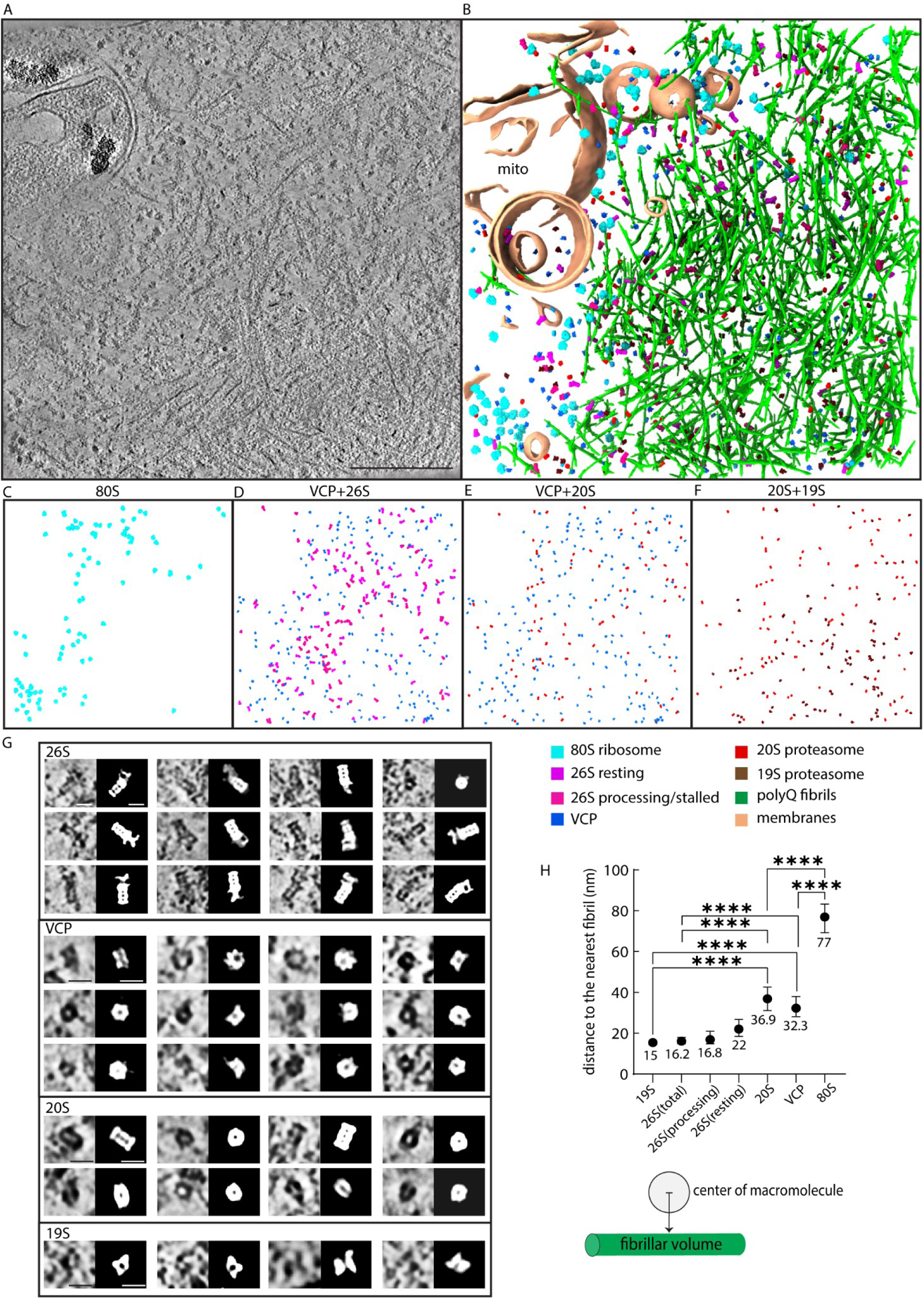
*In situ* visualization of VCP and proteasomes around polyQ aggregates. (**A-F**) A slice of denoised tomogram (**A**) and 3D renderings (**B-F**) illustrating macromolecular distributions around a 150Q aggregate in Neuro2A. **C-F** shows partial renderings of indicated macromolecules. (**G**) Enlarged views of the macromolecules detected in (**A**), along with their coordinates and orientations determined by Stopgap. (**H**) Distance plot of the median distances, with error bars representing 95% confidence interval, measured from the macromolecule center to the nearest fibrillar volume. Schematics of the measurement shown below. Data points collected from 24 tomograms: 19S (n = 2284), 26S total (n = 4400), 26S resting state (n = 1220), 26S processing/stalled state (n = 862), 20S (n = 4880), VCP (n = 4317), 80S ribosome (n = 5420). ****p < 0.0001, Wilcoxon rank test. Scale bars: 250 nm (**A**), 15 nm (**G**).

Denoised ^73^ tomograms with segmentations are displayed to reveal the recruitment of proteasomes and VCP to polyQ aggregates in both Neuro2a (Figure 2A) and HEK293 cells (Figure S3D, S4D). These macromolecules are enriched at the fibrillar periphery, a pattern reflected in their immunofluorescence staining (Figure 1B, 1D). The dense, amorphous polyQ intermediates outside of the fibrillar core upon the inhibition of autophagy ^1^, is mapped with proteasomes and VCP, indicating clear co-localization (Figure S4D-J). Apart from cellular systems where a 20S core typically associates with two 19S regulatory particles as 30S ^40,74^, dividing cells predominantly feature 26S with a 1:1 ratio of 20S to 19S ^75,76^, as is also observed here *in situ*. We classified the 26S into resting and processing/stalled states, distinguished by the distance between the Psmd11 and Psmd12 subunits (Figure S5B). Besides the 26S, we detected a high abundance of 19S and 20S (Figure S5), suggesting that 26S may preferentially dissociate into subcomplexes to sustain protein degradation capacity under proteotoxic stress ^39,66,77^. The 20S core is known to be abundant as a free complex across many cell types^75-77^, and can degrade a variety of disease-associated substrates, including soluble α-synuclein and Tau, which are implicated in Alzheimer’s and Parkinson’s diseases ^78-84^.

From the dataset, we detected VCP in Apo/ADP state or more prevalently, in ATP-bound states (Figure S5A), with the latter exhibiting extra density at one of its N domains (marked by*) not present in the template (Figure S5B). This density may correspond to the UBXL domain of the cofactor (i.e. NPLOC4), while the rest of the density remained flexible and was not resolved. As VCP is known to complex with co-factors NPLOC4 and UFD1 for degrading ubiquitinylated K48-conjugated substrates, we also template matched and detected a class of VCP with extra densities resembling the co-factors (Figure S5B).

We segmented polyQ fibrils and performed nearest distance analysis from the center of macromolecules to the fibrillar volume (Figure 2H). This revealed that the 26S and 19S are closest to the fibrils, suggesting a stronger tendency of being fibril-sequestered, consistent with previous observations of poly-GA ribbons in C9orf72 expansions ^40^. Indeed, aggregated polyQ is well-documented to stall the UPS ^42,43,85,86^, while more soluble polyQ can be degraded ^44^. The analysis showed that VCP and the 20S were farther from the fibrils, with median distances exceeding 30 nm, yet still closer than the 80S ribosomes, which had a median distance of over 70 nm. Since 80S ribosomes are not known to interact with polyQ fibrils, their distance serves as a non-binding control in the pairwise distance measurements. In line with the mass spectrometry analysis (Figure S1A-E), 20S is less sequestered by the aggregated fibrils in comparison to the 19S, and likely associates more with the polyQ intermediates at the fibrillar periphery.

### 4. *In situ* macromolecular sociology suggests cooperativity between VCP and the proteasomes

We reasoned that collecting a dataset with a smaller pixel size can improve resolution for the reconstructions; therefore, 410 tomograms were acquired at 1.89 Å using a more advanced setup (Figure 3, Figure S6-8) to optimize signal detection for smaller complexes, including the 20S, 19S, and VCP, at 750, 850, and 540 kD, respectively. The resulting coordinates were mapped back onto the tomographic volume for inspection (Figure 3A-D, S6D). For improved resolution in the final reconstructions, we used the Warp-Relion-M workflow ^87,88^, achieving near Nyquist resolution for binned subtomograms of 80S ribosomes as a benchmark (Figure S6E, S7A).

**Figure 3.**
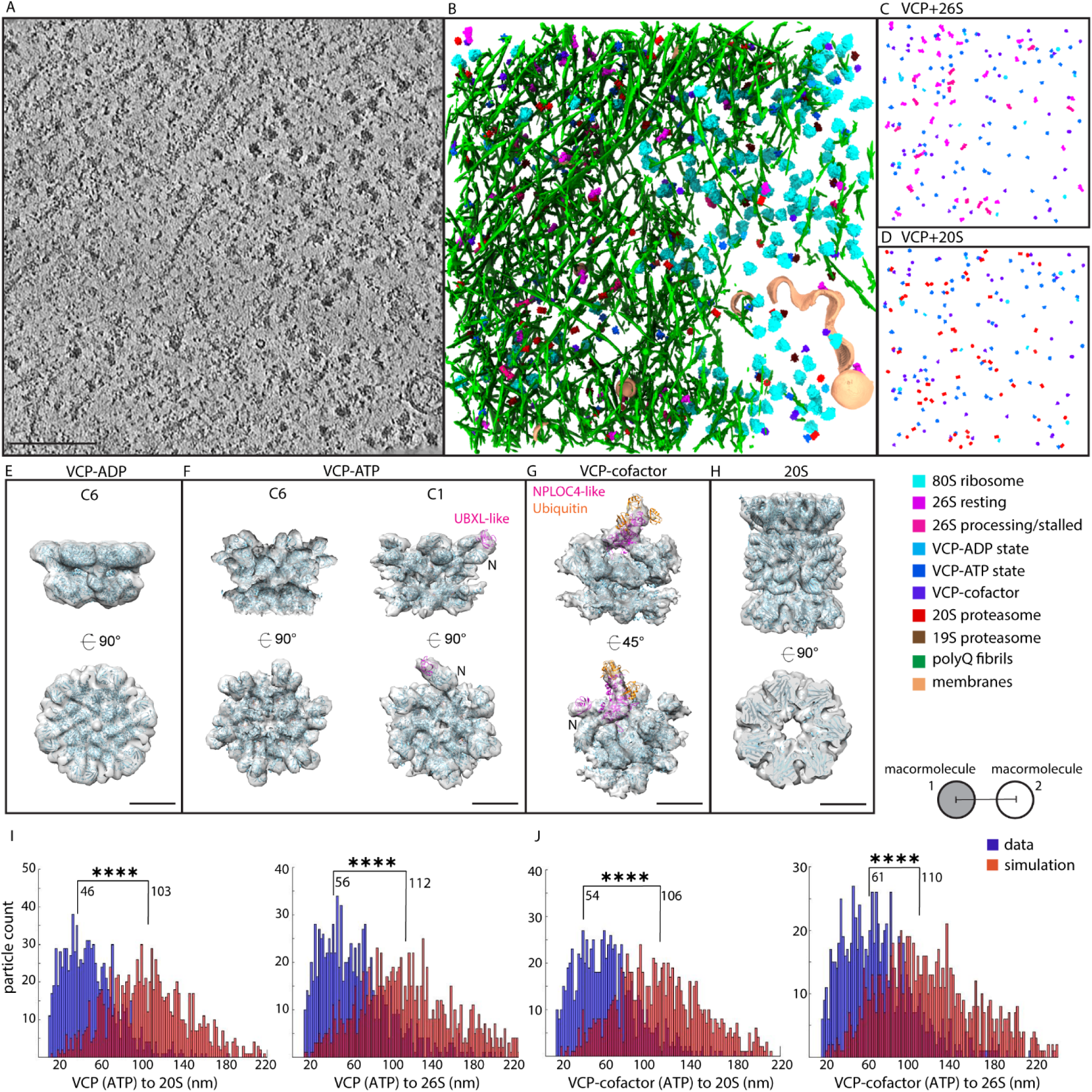
*In situ* macromolecular sociology suggests cooperativity between VCP and the proteasomes. **(A-D)** A slice denoised tomogram (**A**) and the 3D renderings (**B-D**) of a 150Q aggregate illustrating macromolecular distributions. (**C-D)** shows partial rendering of macromolecules. **(E-H)** *In situ* reconstructions of VCP and 20S docked with PDB structures in ribbon representation. The VCP-apo/ADP state is docked with PDB 5FTK (**E**), VCP-ATP state is docked with PDB 7JY5 and NPLOC4 UBXL domain (**F**), and the VCP-cofactor complex is docked with PDB 7LN3, NPLOC4, and 3 ubiquitins (**G**). The N domain of VCP is labeled (**F, G**). The 20S is docked with PDB 7V5M (**H**). (**I-J)** Nearest distance measurements between particle centers for VCP and the 20S or 26S proteasomes, compared to random simulations (pink), with a minimum cutoff of >10 nm. Median distance values are displayed, ****p < 0.001 Kolmogorov-Smirnov test, n =15 tomograms. Schematics of the measurement shown to the right. Scale bar: 150 nm (**A**), 5 nm (**E-H**).

The VCP reconstructions confirmed the observation of VCP in the ADP/apo state, the ATP-bound state without and with the extra NPLOC4-like cofactor density in a ∼3:20:20 ratio (Figure 3E-G, S7B-C). The reconstruction of the VCP ATP-bound state with C6 symmetry was docked with PDB 7JY5 (Figure 3F). As VCP is known to adopt multiple states that lack symmetry ^89^, we also performed reconstruction without symmetry constraints, confirming the observation of the UBXL domain-like density on one of the N domains (Figure 3F, S7B). In the VCP-cofactor co-complex, where the rest of NPLOC4-like density is resolved (Figure 3G, S7C), three ubiquitin appeared to be fitted into the density. The co-complex is weakly connected between the D1 and D2 ATPase rings, positioned at an angle that is better docked with PDB 7LN3 (Figure 3G), suggesting substrate translocation and processing in asynchronous ATP hydrolysis ^89^.

The 20S reached an overall resolution of 10 Å with D7 symmetry imposed, about 10% of the particles were classified to have a different gate configuration, a state that may indicate higher substrate engagement (Figure 3H, S8A). The 26S were classified into resting and processing/stalled states (Figure S8B), and due to the flexibility of the 19S, the resolution was lower, revealing only the resting state (Figure S8C).

Using dataset 2 (Figure 3I-J) and dataset 1 (Figure S9), we conducted nearest distance analyses to explore the spatial relationships among VCP, 20S, and 26S. As negative controls, simulations were performed with randomized coordinates in the same volume, excluding areas occupied by organelles, membranes, and fibrils (Figure S9A). Both datasets showed a significantly closer proximity between centers of the macromolecules compared to the random simulation. Besides the anticipated VCP-26S cooperativity ^51^, VCP is also closely associated with the 20S with median values at 46-54 nm (Figure 3I-J), a pattern that is conserved across mouse and human cells (Figure S9B-C). The proximity between VCP and the 20S suggests a direct functional relationship, potentially reflecting VCP’s cooperativity with the 20S catalytic core in a role akin to the well-characterized 19S proteasome ^48^. However, the variability in the VCP-20S distance suggests that the docking of VCP with the 20S as a stable co-complex is unlikely, pointing instead to a more transient cross-talk.

### 5. VCP and 20S cooperativity explained by the VCP C-terminal HbYX motif

Our *in situ* nearest distance analysis suggests that VCP and the 20S cooperate in polyQ degradation. To investigate this, we conducted *in vitro* degradation assays with purified human 20S and VCP complexes (Figure 4A-D, S10A-B). Since aggregated 64Q cannot be cleared by VCP or the proteasome (Figure 1A), the more soluble 64Q fraction was used as the substrate, upon isolation from Neuro2a cells using GFP-trap pull-down followed by the removal of large inclusions with a filter (0.2 µm). 64Q was incubated with either the 20S alone or in combination with VCP and/or the 19S. 20S alone had minimal effect on substrate degradation over the tracked time frame (Figure 4A). When VCP and cofactors NPLOC4/UFD1 (UN) were included, an enhancement in polyQ degradation was observed (Figure 4B). The combined effect of 19S and VCP was more pronounced, at the 2 h time point (Figure 4C-D).

**Figure 4.**
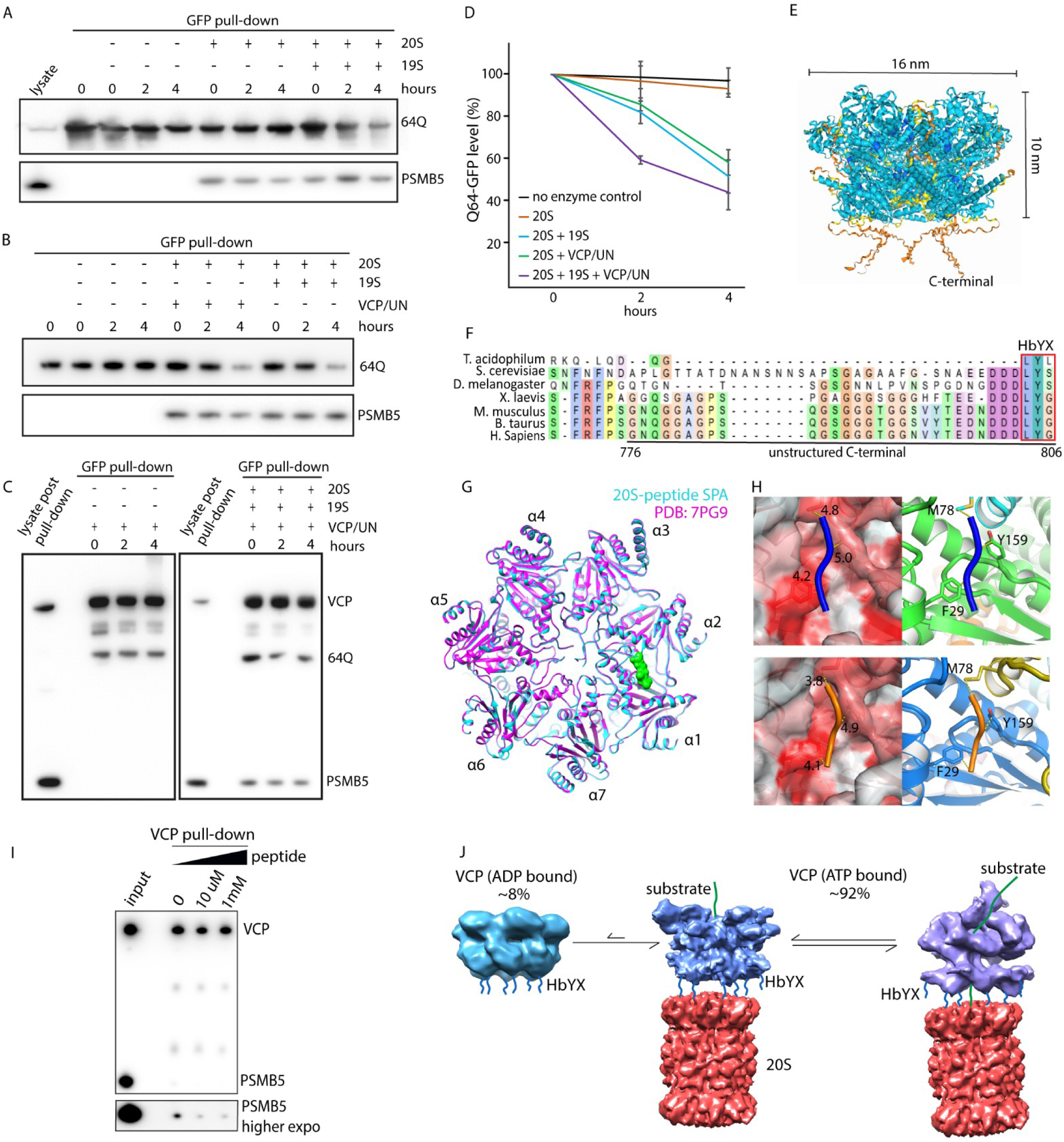
VCP and 20S cooperativity explained by the VCP C-terminal HbYX motif. (**A-C**) *in vitro* degradation assay with GFP-trap pull-down for non-aggregated 64Q, followed by incubation with indicated enzymes (2 μg) for the indicated time. VCP cofactors NPLOC4/UFD1 referred to as UN. Immunoblots probed with the indicated antibodies, 64Q blotted with antibody 3B5H10. (**D**) Plot summarizing 64Q degradation pattern, n = 3, error bar: s.d. (**E**) Alphafold3 prediction of the human VCP hexamer, including its disordered C-terminal tail. (**F**) Sequence conservation of VCP or homologs across archaea and eukarya, with the C-terminal HbYX motif boxed. (**G**) The VCP C-terminal peptide, traced with poly-alanine (green surface view), is positioned between the PSMA1 and PSMA2 subunits in complex with the human 20S (cyan) from cryo-EM SPA, overlaid with PDB 7PG9 (magenta, RMSD: 0.17 Å). (**H**) Modeling of the 20S-peptide binding pocket from cryo-EM SPA reveals four peptide residues positioned within the hydrophobic pocket between PSMA1 and PSMA2. Panels highlight hydrophobic residues in red (left) and label them (right): Phe29 and Tyr159 (PSMA1), and Met78 (PSMA2). The distances from peptide backbone to the hydrophobic residues are labeled. (**I**) VCP-20S binding assay with peptide added at increasing concentrations, followed by VCP pull-down for 20S, blotted with the indicated antibodies. (**J**) Model: macromolecular sociology of VCP and the 20S in processing polyQ intermediates *in situ*, with the abundance of the VCP states approximated from the *in situ* cryo-ET data.

Each of the six VCP/p97 subunits possesses an unstructured C-terminal tail that terminates in a highly conserved HbYX motif (Figure 4E-F). Spanning residues 776-806, this tail can extend several nanometers in length. We hypothesize that the VCP HbYX motif, similar to those found in the 19S or PA200 complexes ^48,90^, bind to the 20S in the inter-subunit pockets of PSMA and facilitate the recruitment of VCP. We thus incubated the human 20S with a peptide containing the VCP C-terminal HbYX motif (DNDDDLYG) for cryo-EM SPA ^91^. Our results showed that 35% of 20S particles resembled PDB 7PG9 at 3.7 Å with no bound peptide (Figure S10C-D). Another 25% (4.3 Å) exhibited poorly resolved densities (>7 Å) in multiple inter-subunit pockets of PSMA (Figure S10E). And 16% of h20S displayed a strong signal in the pocket between PSMA1and PSMA2 at a global resolution of 3.9 Å (Figure 4G-H, S10F-G). Despite the small size of the peptide, the density extends across the pocket and interacts with both PSMA1 and PSMA2, primarily through hydrophobic interactions, as observed in the archaeal HbYX-20S co-complex ^91^ (Figure 4H). The distances from the peptide backbone to Phe29 and Tyr159 of PSMA1 and Met78 of PSMA2 are in 3.8-5.0 Å range. Consistent with the SPA result, AlphaFold3 ^92^ predicts that all seven pockets between the human 20S α subunits can accommodate such a motif (Figure S10H). Furthermore, *in vitro* binding assay performed with purified VCP and 20S showed that their binding can be prevented with increasing concentrations of free VCP HbYX peptides (Figure 4I).

In comparison to the free human 20S, the binding of the VCP HbYX motifs from single-particle analysis (Figure 4G, S10E-F) or in the Alphafold3-predicted structure (Figure S10H) did not show a significant shift in α ring for gate expansion, as observed in the archaeal 20S ^91,93^. The lack of clear gate expansion in human 20S was also evident when overlaying the free and processing states of 20S (PDB 7PG9 vs. 7V5M/7V5G) ^77^(Figure S10I). Overall, our study supports a model in which VCP cooperates with the 20S through flexible interactions, facilitating the coupling of VCP-mediated substrate unfolding with degradation by the 20S catalytic core (Figure 4J).

## Discussions

Building on our previous *in situ* cryo-ET work revealing the preferential clearance of non-fibrillar polyQ by the macroautophagy machinery ^1^, we now show that VCP and proteasomes of the UPS pathway also target the more soluble amorphous polyQ at the fibrillar periphery. The relevance of VCP in polyQ clearance has been demonstrated previously, as overexpression of VCP or its cofactors can mitigate polyQ toxicity in both yeast and neurons ^25,65^. In addition to being recruited by ubiquitinylated polyQ, the recruitment of VCP and proteasomes is known to be facilitated by induced ER stress and associated degradation ^25,65,94,95^. The yeast homolog of human VCP, Cdc48, along with its cofactor NPLOC4/UFD1, was first identified to facilitate the extraction and unfolding of ubiquitinated proteins from the ER^94^. Our findings suggest that activating the ER quality control system to simultaneously recruit VCP and proteasomes that otherwise accumulate in the nucleus (Figure 1B, 1D), may enhance the removal of the proximal soluble polyQ intermediates -- known to be the most toxic form contributing to disease progression ^14,18,24-27^.

We elucidated the *in situ* structures of VCP and the proteasome in multiple states, uncovering an unexpected proximity between VCP and the 20S (Figure 4J), in addition to the anticipated VCP-26S cross-talk. Compared to neurons ^40^, the density of the 26S around the disease aggregate was lower than that of the 19S and 20S subcomplexes in dividing cells, making the stalled 26S less visually pronounced alongside the fibrils. However, the tendency of fibrillar polyQ to recruit the 26S proteasome or its 19S subcomplex appears conserved, given their close proximity (Figure 2H). It is conceivable that Neuro2A and HEK293 cells have higher expression of proteasome regulators (i.e. Ecm29) ^39,66-68^, which facilitate the dissociation of 26S into 19S and 20S under proteolytic stress, thereby preserving 20S core functionality.

The *in situ* proximity between VCP and the 20S is likely mediated by VCP’s C-terminal HbYX motif, which can bind into the pockets between the PSMA subunits. This is supported by our 20S-HbYX peptide co-complex structure from cryo-EM SPA (Figure 4) that aligns with Alphafold3 predictions. Together these observations indicate that VCP may function similarly to the 19S in facilitating the unfolding of polyQ intermediates for coupled degradation by the 20S. While evidence from archaea indicates that VCP stably binds to and activates the 20S to enhance substrate degradation, similar evidence in higher eukaryotes remains limited ^96-98^. While it is known that the gate conformation at the two opposite ends of the 20S proteasome are coupled, the binding of one regulatory ATPase may allosterically open the opposite gate ^99^. This may explain why VCP-26S demonstrates higher degradation activity at 2 h (Figure 4D), potentially aided through VCP’s direct binding to the 20S barrel on the opposite side of the 19S.

The *in situ* proximity of VCP to the 20S proteasome observed in our study supports the idea that promoting 20S activity, facilitating gate opening, or enhancing VCP-20S coupling could be promising strategies for treating amyloid-related diseases. Supporting this, *C elegans* with hyperactive 20S gate-open mutant exhibit increased lifespan and resistance to proteotoxic stress ^100^. Conversely, hyperactive VCP mutants (A232E, R155H) cause neurodegeneration ^59-62^, likely due to VCP unfolding ubiquitinylated substrates that are not immediately degraded by the proteasome, making them prone to aggregation and propagation ^51^. In this context, VCP and its K48 ubiquitin-specific cofactors (NPLOC4/UFD1) may be viewed as competitors of the 19S for ubiquitinylated substrates, highlighting the necessity of maintaining robust VCP-proteasome cross-talk. Our findings suggest that stabilizing VCP-20S coupling could provide an alternative pathway, bypassing reliance on 19S subcomplexes, which we show (Figure S1A-C, Figure 2H) to be more avidly sequestered by fibrillar polyQ aggregates.

## Limitations

One technical limitation is that the 2-day induction of the shorter 64Q-GFP primarily results in soluble oligomeric forms without significant ER enrichment; therefore, pharmacological induction of ER associated degradation did not produce a discernible phenotype. A second limitation is that the 2-day induction of the longer 150Q-GFP leads to predominantly SDS-insoluble aggregates, which obstruct sample loading wells during immunoblotting and clog the filter trap device in slot blot assays, preventing accurate quantification by blotting.

While VCP is known to be involved in the activation of autophagy ^101^, our cryo-ET analysis did not reveal a high abundance of autophagy-related structures without pharmacological induction. Therefore, our *in situ* data do not address VCP’s potential contributions to autophagy.

## Method details

### Cell lines, Plasmids, siRNAs, CRISPR/Cas9 knockout, and Chemicals

Cells were grown with DMEM + Glutamax (Gibco) with 10% FBS (Gibco) and 100 units/ml Penicillin-Streptomycin (Gibco) in incubator at 37°C with 5% CO2; no unusual Hoechst staining (Cell signaling) observed for mycoplasma contamination. For passages, cells were washed in PBS (Gibco), trypsinized with TrypLE™ Express (Gibco). Neuro2a containing stable expression of the inducible 64Q- or 150Q-GFP were described before ^14^. For the induction of the transgene expressing polyQ, muristerone A (Abcam) was applied at 1 μM for 2 days. The following chemicals were applied to cell culture: tunicamycin (Sigma), MG132 (Sigma), and NMS873 (Sigma).

For transient expression of 97Q-GFP, HEK293 were transfected for 48 h using Lipofectamine 3000 (Invitrogen) in Opti-MEM (Gibco) as per manufacturer’s protocol, media was refreshed after 1 day. The plasmids expressing GFP tagged Htt97Q exon 1 were previously described ^41,102^.

### Immunoblotting and filter trap assay on cellular lysates

Cells expressing 64Q-GFP from 12-well plates were lysed on ice in 80 µL RIPA buffer (Thermo) supplemented with protease inhibitor cocktail (Roche) and Benzonase (Novagen) with intermittent vortexing. Protein concentration in total cell lysates was determined using Protein Assay Dye Reagent Concentrate (Bio-Rad), normalized before denaturing in 4x LDS sample buffer (Thermo) containing 2.5% β-mercaptoethanol (Sigma) and boiling at 95°C for 5 min. Proteins were separated on NuPAGE 4-12% bis-tris gels (Thermo) with MES running buffer (Thermo). Afterwards, proteins were transferred to PVDF membranes (Roche) in tris-glycine buffer using semi-dry transfer. Membranes were washed in TBS-T and blocked in 5% low-fat dry milk (Sigma) dissolved in TBS-T for 1 hour at room temperature. Blots were incubated with primary antibodies (1:500-1:1000) overnight at 4°C, washed three times with TBS-T and probed with HRP-conjugated secondary antibodies (1:10000) for 1-2 h at room temperature. Chemiluminescence was developed using HRP substrate Immobilon Classico (Merck) or SuperSignal West Dura (Thermo), on an ImageQuant800 (Amersham) imager with control software v1.2.0. Intensity of protein bands was quantified using Fiji/ImageJ ^103^.

Filter-trap assay for the detection of aggregated 64Q-GFP was performed with lysates prepared as above. 10 or 20μg total protein was diluted in 100 - 200μL RIPA. Cellulose acetate membrane (0.2 μm pore, GE Healthcare) was equilibrated in 0.1% SDS/H_2_O and fixed to the filter trap device (PR648 Slot Blot Blotting Manifold, Hoefer). Samples were loaded under vacuum. Slots were then washed with 200 μl 0.1% SDS/H_2_O three times followed by standard immunoblotting.

### 150Q-GFP pull-down for label free quantitative mass spectrometry

Htt150Q were induced with muristerone A for two days in 15 cm plates, and was lysed on ice in buffer (20mM Tris pH 7.4, 100mM NaCl, 0.5% NP40) with protease inhibitor cocktail and benzonase, lysates were passed multiple times through 23-gauge needle to enhance lysis. The lysate was pelleted at 1000g at 4C for 10 min to remove large cellular clumps. To partition and collect for the aggregated Htt150Q, the lysate was pushed through pre-equilibrated 0.2 μm PVDF syringe filter (Millipore), and the trapped fractions were collected from the filter in the same buffer. Next, the aggregated fraction was incubated with GFP-Trap and control beads (ChromoTek) at 4°C for 6 h. The beads were washed in the same buffer, and then analyzed by label free LC-MS/MS. For the analysis of the total Htt150Q with GFP-Trap pull down, a similar protocol was followed but without the step of passing and collection from the 0.2 μm filter.

For LC-MS/MS, proteins were reduced and alkylated in SDC buffer (1% Sodium deoxycholate, 40 mM 2-Cloroacetamide (Sigma), 10 mM TCEP (Thermo) in 100 mM Tris pH 8.0) for 20 min at 37 °C. The samples were then diluted with MS grade water (VWR) and digested overnight at 37 °C with 1 µg Lys-C (Labchem-wako) and 2µg trypsin (Promega), followed by acidification with Trifluoroacetic acid (Merck) to 1% (pH < 2). Next, the samples were purified via Sep-Pak Vac 1cc (50mg) tC18 Cartridges (Waters GmbH) with 0.1M acetic acid (Roth) wash and eluted with 80% Acetonitrile and 20mM acetic acid (Roth), and vacuum dried for resuspension in Buffer A (0.1% (v/v) Formic acid (Roth)). Peptides were then loaded onto a 30-cm column (packed with ReproSil-Pur C18-AQ 1.9-micron beads, Dr. Maisch GmbH) via the Thermo Easy-nLC 1200 autosampler (Thermo) at 60°C. Using the nano-electrospray interface, peptides were sprayed onto the Orbitrap MS Q Exactive HF (Thermo) in buffer A at 250 nl/min, and buffer B (80% Acetonitril, 0.1% Formic acid) was ramped to 30% in 60 min, 60% in 15 min, 95% in 5 min, and finally maintained at 95% for 5 min. MS was operated in a data-dependent mode with survey scans 300 - 1650 m/z (resolution of 60000 at m/z =200), and up to 10 top precursors were selected and fragmented using higher energy collisional dissociation (HCD with a normalized collision energy value at 28). The MS2 spectra were recorded at a resolution of 30000 (at m/z = 200). AGC target for MS and MS2 scans were set to 3E6 and 1E5 respectively within a maximum injection time of 100 and 60 ms for MS and MS2 scans respectively. Dynamic exclusion was set to 30ms.

Raw MS data were processed using the MaxQuant platform ^104^ with standard settings, and searched against the reviewed Human or Mouse Uniprot databases, as well as polyQ-GFP sequences, allowing precursor mass deviation of 4.5 ppm and fragment mass deviation of 20 ppm. MaxQuant by default enables individual peptide mass tolerances. Cys carbamidomethylation was set as static, and Ub, Met oxidation and N-terminal acetylation as variable modifications. Protein abundances within a sample were calculated using iBAQ intensities ^105^ and were quantified over the samples using the LFQ algorithm ^106^ for analysis with the Perseus software (https://maxquant.net/perseus/).

### Confocal fluorescence light microscope image acquisition and analysis

Cells were seeded onto glass coverslips in 6-well cell culture plates. Following growth and treatment, cells were washed with PBS and fixed in 4% paraformaldehyde (Sigma) for 10 min, permeabilized with 0.1% Triton X-100 (Sigma) for 5 min, and blocked in 5% milk dissolved in PBS at room temperature for 1 h. Primary antibodies were applied at a dilution of 1:50 - 1:100 in block overnight at 4°C, then washed three times in PBS (5 min) and incubated with secondary antibodies at a dilution of 1:1000 in dark at room temperature for 1 h. Coverslips were washed three times with PBS (5 min), mounted onto glass slides with Fluoromount (Sigma).

For staining with ER-Tracker or LysoTracker Red DND-99 (Invitrogen), the dyes were added to the culture prior to fixation or vitrification according to the manufacturer’s protocol. The inclusion of Lysotracker stain limits the downstream analysis to cytosolic polyQ aggregates. 30 min prior to vitrification or live confocal light microscope imaging, cells were stained with Annexin V-pacific blue (Invitrogen) or iFlour 555 (Abcam) according to the manufacturer’s protocol to quantify and exclude apoptotic cells from further analysis.

Fluorescence microscopy data were acquired at the MPI imaging facility, on a confocal laser scanning microscope (Zeiss LSM 980, Jena, Germany) with a PLAN/APO 63x oil objective and the Zeiss Zen software. Images for quantifications of control and the treatments were taken with the same exposure setting, data analysis was performed in Fiji/ImageJ. Representative images and datapoints from reproducible experiments are shown. The measurements of the aggregate cross-sectional area were performed with the central zone thresholded to > 8000 for GFP intensity (threshold tool), and the peripheral region threshold to 800-8000 for the intensity. Data quantifications were performed in GraphPad Prism 9 (Graphstats Technologies), p values generated from two-tailed student’s t-test.

### Sample vitrification for cryo-ET

Cells were seeded on holey carbon-coated 200 mesh gold R2/1 EM grids (Quantifoil Micro Tools, Jena, Germany) in a 35 mm cell culture dish (MatTek). Prior to vitrification, cells were applied with DMEM containing 10% glycerol as a cryo-protectant and Dynabeads (Invitrogen) at 1:40 dilution for 3D CLEM workflow, then immediately mounted on Vitrobot Mark IV (Thermo), blotted from the back side using FEI Vitrobot Perforated Filter Paper (Whatman) with force 10 for 15 seconds at room temperature and humidity, and plunged into a 2:1 ethane:propane mixture cooled down to liquid nitrogen temperature. Plunge-frozen grids were then clipped into Autogrid support frames modified with a cut-out (Thermo), stored in liquid nitrogen, and maintained at ≤ −170°C for all steps.

### Correlated light-electron microscopy (CLEM) and cryo-focused ion beam (FIB) milling

Vitrified sample on autogrids were loaded onto a cryo-confocal set up (Leica SP8) equipped with a 50X/0.9 NA objective (Leica Objective), metal halide light source (EL6000), air-cooled detector (DFC900GT), a cryo-stage (-195°C), and two HyD detectors. The sample was kept in liquid nitrogen vapor, following a similar workflow as described ^71^. Cryo-confocal z-stacks (step size 500 nm, x-y pixel size 85 nm) were taken with pin hole = 1, and a 7 μm depth with the LAS X Navigator software, using 488 and 552 nm laser excitation for GFP tagged proteins and LysoTracker Red, respectively. To improve signal clarity, image stacks were de-convoluted and restored with Huygens Essential software (Scientific Volume Imaging) to remove noise. The stacks were then imported into the 3Dcorrelation software ^69^ and re-sliced into cubic voxels.

For preparing the lamella (150-250 nm), autogrids were mounted into an Aquilos II FIB/SEM system (Thermo) at < −180 °C. To protect the milling front of the lamella, gaseous organometallic platinum was sprayed onto the sample on the cryo-stage using a gas injection system. To target the cell for milling, the grid square was correlated with the cryo-confocal fluorescence z-stack using 3Dcorrelation ^69^. The target was imaged and correlated iteratively with the z-stack throughout milling for accuracy. The 12-15 μm wide lamellas were generated using a Gallium FIB at 30 kV with a 20° stage angle in three consecutive steps. The more distant region (> 2 μm) above and below the target was rough milled with a high current of 500 pA, followed by fine milling to a ∼800 nm using 100 pA. A final polishing of the lamella to 150-250 nm was carried out with 30-50 pA current. Lamella final thickness was estimated with SEM at 3 keV ^107^, for the lack of overcharging.

### Cryo-ET data acquisition, tomogram reconstruction

For dataset 1, 250 tilt series were collected on a Titan Krios cryo-TEM at 300kV with a field emission gun, a post column energy filter (Gatan, Pleasanton, CA, USA) operating at zero-loss with a 20eV slit width, and a K2 summit direct director (Gatan). The energy filter was used to increase image contrast. Low-magnification were taken at 8700x (object pixel size 1.69 nm) to obtain lamella overviews, it provided enough detail to locate the aggregates and surrounding structures for data acquisitions. High-magnification (42,000x at 3.35 Å pixel size) tilt series were collected at sites of interest using SerialEM ^108^, operating in low dose mode with tracking and focus enabled. Tilt series were taken with a 2° tilt increment with an angular range from ∼−70° to 50° starting at -10° to compensate for the lamella pre-tilt, with dose-symmetric tilt scheme ^109^. The K2 camera operating in dose fractionation counting mode, recorded frames every 0.2 s for 2 electrons/Å² per tilt. The cumulative dose was in the range of 80-120 electrons/Å² per tilt series. Targeted defocus of -5 to -3 μm were applied to boost contrast. FIB-related imperfections were removed for the lamella overviews, using the LisC filter algorithm ^110^.

For dataset 2, 410 tilt series were collected on a Titan Krios G4 instrument at 300 kV equipped with a Selectris X energy filter operating at zero-loss with a slit width of 10eV, and a Falcon 4i camera (Thermo). Low-magnification were taken at 11500x (object pixel size 2.13 nm) to generate lamella overviews. Tilt series were recorded at 1.89 Å pixel size using the Tomography 5 (Thermo) in EER file format, with a dose-symmetric tilt scheme. The tilt series were collected with a 3° tilt increment with an angular range of ∼120°, and with cumulative dose of 80-120 electrons/Å². Targeted defocus was in the range of -5 to -3 μm.

Raw frames were pre-processed using in-house Matlab ^111^ wrapper scripts (Tomoman: https://github.com/williamnwan/TOMOMAN) ^112^. The relative shifts of the image between camera frames due to stage drift and beam-induced motion were corrected by MotionCor2 ^113^, followed by exposure filtering ^114^. The EER frames were motion corrected using Relion’s implementation of MotionCor2 ^115^ followed by exposure filtering. Bad tilts were removed upon manual inspection using the Tomoman script. The tilt series were then aligned using patch tracking, binned, and reconstructed by weighted back projection in IMOD ^116^. CTF was estimated by tiltCTF ^117^ and the final CTF corrected reconstruction was made in IMOD for template matching. For figure display, tomogram contrast was enhanced upon denoising with cryoCARE, by which tilt series were separated into odd and even tilts during motion correction^73^.

### Cryo-ET template matching

The real-space noise-correlation-based template-matching approach implemented in STOPGAP 0.7.0 ^72^ was performed. Template matching was carried out with the dataset 1 (pixel size 3.35 Å) by four-fold binning to a pixel size 13.4 Å, and repeated with dataset 2 (pixel size 1.89 Å) by eight-fold binning to a pixel size of 15.12 Å. For this work, the parameters were set as follows: 15° angular scanning step, low-pass filter radius = 10, high-pass filter radius = 1, apply_laplacian = 0, noise_corelation = 1 and calc_ctf = 1. Low-pass filtered PDB structures (40 Å) were generated using the molmap command in Chimera ^118^ as initial models for template matching, with masks applied. Masks for the template matching were created using relion_mask_create with extend_inimask = 3 and width_soft_edge = 3. The above setting was optimized to produce distinguished peaks for visualization in IMOD ^116^.

To mitigate against template bias, several validation tests were conducted: 1) Low-pass filtered template were truncated to miss part of the 26S. Template matching using test dataset allowed the recovery of the full structure. 2) Template matching for the 20S proteasome and VCP was performed with cylindrical volumes, the recovered particles converged to the expected structures. 3) low-pass filtered hybrid structures, averaging known states from PDB models (5FTK, 5FTN) were used as templates, enabling the recovery of multiple states of VCP.

Upon template matching, the cross-correlation cut-off value was determined separately for each tomogram by visually inspecting the hits overlaid on the tomogram in IMOD. To reduce false positives, a mask was created in Amira (Thermo) to exclude regions above and below the lamella, at tomogram edges, and organelles along membranes traced with Membrain ^119^. The subtraction of membranes from the tomograms prevents the selection of membrane-bound macromolecules for downstream analysis. To analyze all potential macromolecules of interest in the datasets, oversampling in the initial stage was intended to lower false negative assignments. The coordinates of the macromolecules were extracted as subtomograms in STOPGAP.

### Cryo-ET subtomogram averaging

Iterative 3D classifications, 3D alignments, and repeated template matching on the complete dataset, using data-derived reconstructions as templates, together, improved the final resolution of the reconstructions. 3D classifications were performed using simulated annealing stochastic hill-climbing multireference alignment in STOPGAP ^72^. 3D classifications for the 80S ribosomes and 26S proteasomes were initially performed with four-fold binned (13.4 Å) and eight-fold binned (15.12 Å) subtomograms for dataset 1 and 2, respectively. For the smaller macromolecules (VCP, 20S, and 19S), initial 3D classifications were carried out with two-fold and four-fold binned subtomograms, for dataset 1 and 2, respectively. Initial classifications typically began with 8-12 classes, using *de novo* references generated in STOPGAP using random particle initial sets with global circular masks applied. 3D classification parameters (search_mode = shc, temperature = 10 to 0, iterations = 25 to 30, angincr = 0, angiter = 0, phi_angincr = 2, phi_angiter = 2) were used, with D7 symmetry imposed for 20S, C6 or C1 symmetry for VCP, and C1 symmetry for the rest of macromolecules. Class convergence during iteration cycles was monitored using STOPGAP scripts (sg_motl_plot_class_bar_graph.m, sg_motl_plot_class_changes.m, sg_motl_plot_class_convergence.m) to assess completion.

Once well-aligned particle classes were obtained, as assessed by the reconstruction and corresponding gold-standard FSC curve (cut-off of 0.143) of the two independent halfsets, selected classes were re-extracted at a smaller binning for iterative alignment and classifications. The final coordinates of macromolecules (26S, 20S, 19S, VCP, and 80S ribosomes) were pooled, further refined using the script (sg_motl_distance_clean_xyz.m) by ensuring that adjacent macromolecules from all types were no closer than 10 nm center-to-center. In cases of any conflict, only the macromolecule with the higher cross-correlation score was retained. Conflicts occurred in 10% of macromolecules from dataset 1 and 5% from dataset 2, suggesting that iterative STOPGAP classifications effective in reducing false-positive assignments.

For estimating the final resolutions of the reconstructions in Relion, the cleaned coordinates of a particle type were imported into Warp 1.0.9 ^87^ for reprocessing, which filters out tomograms and particles with abnormal CTF and astigmatism. The resulting particles were extracted two-fold binned in Warp for 3D refinement in Relion 3.0 ^120^, first globally and then locally with a sigma of 0.5 and search angle of 0.9-1.8°. Finally, particles were imported into M 1.0.9 ^88^ for further refinement of geometric and CTF parameters to improve resolution, which proved highly effective for larger macromolecules (i.e. 80S ribosomes). Final visualization of the reconstructions was conducted using UCSF Chimera for surface views ^118^and IMOD for XYZ views. Fibril and membrane segmentations, along with macromolecules, were displayed in ChimeraX for visualization ^121^. The number of subtomograms and resolution estimates for different macromolecules obtained from template matching and subtomogram averaging for datasets 1 and 2 are documented in Figure S5A and S6E, respectively.

### Nearest distance analyses

Nearest pairwise distance calculations were performed on tomograms capturing the peripheral region around polyQ fibrils, where the macromolecules of interest were densely distributed. Relatively thinner tomograms with sufficiently high particle cross-correlation scores were selected to ensure that detection was not limited by a low signal-to-noise ratio. The analysis was based on previously developed algorithm ^122^. For the simulation, the script creatSimuStar.m first placed the first macromolecule type into the tomographic volume based on its coordinate file. It then applied masks to exclude regions occupied by fibrils, organelles, and area outside the lamella plane, thereby defining the space where the second macromolecule type can be randomly placed. The script then generated a simulated coordinate file and a volume file, to which binary threshold -- derived from the reconstruction and estimated in Chimera – was applied. Subsequently, the script mainAnalysis.m calculated and plotted the shortest neighboring distances between the centers of the two macromolecule types in both real data and the simulation. A Kolmogorov-Smirnov statistical test was performed to assess the significance of the distance distributions.

Nearest distance analysis between macromolecule centers and the fibrillar volumes was performed with the polyQ fibrils, detected as previously described with the XTracing module ^123^, segmented in Amira. Briefly, tomograms were denoised using a non-local means filter, and fibril-containing regions were identified by searching for a cylindrical template with an 8 nm diameter and 42 nm length. Once fibrils were reliably detected and traced in Amira, a segmentation volume file was generated. Euclidean distances were then converted to physical units based on the pixel size, allowing for the calculation of macromolecule-to-fibril distances. Wilcoxon rank test was applied to assess the significance of the distance distributions.

### Single particle analysis

The H-DNDDDLYG-OH peptide was synthesized at the MPI Biochemistry core. For single particle analysis, 20S (1 μg/ul) was incubated with 1mM DNDDDLYG peptide dissolved in PBS, for 15 min in room temperature ^91^. Copper EM grids with 200 mesh R1/4 carbon film (Quantifoil Micro Tools, Jena, Germany) was glow-discharged, followed by sample vitrification on a Vitrobot Mark IV (Thermo) at 4°C and 100% humidity. 3 μl of the sample was applied, blotted using FEI Vitrobot Perforated Filter Paper (Whatman) with force 3.5 for 4 second.

The vitrified sample was collected on a Titan Krios cryo-TEM operating at 300 kV with a field emission gun, a post-column energy filter (Gatan, Pleasanton, CA, USA) set to zero-loss mode with a 20 eV slit width, and a K2 Summit direct detector (Gatan). Data were acquired at a pixel size of 1.69 Å, targeted defocus range of -0.5 to -3 μm, with 5552 movies using SerialEM and processed in Relion 4.0 ^124^. After 2D and 3D classifications to clean the dataset, > 300K 20S particles remained. Local 3D classification without alignment, with a mask around the 20S alpha subunits led to a few classes, with 35% particles containing no additional density in comparison to known human 20S (PDB 7PG9), and 25% particles containing multiple weak densities, and 16% of the particles containing density between PSMA1 and PSMA2. Postprocessing was performed with B factor = -120 applied.

For model building, the 20S proteasome (PDB 7V5M) ^77^ was fitted to the 20S-peptide co-complex using Chimera ^118^, and poly-Ala backbones of peptide DDDLYG were manually built in COOT ^125^. The entire model was refined using phenix.real_space_refinement ^126^. The hydrophobic surface surrounding the bound peptide was visualized in PYMOL (Schrödinger Inc), and all figures were prepared using PYMOL.

### Protein purification and ATPase activity assay for VCP and co-factors NPLOC4/UFD1

Full-length human VCP, UFD1, and NPLOC4 were overexpressed in *E. coli* Rosetta (DE3) pLysS cells. Cells expressing 6×His-tagged VCP were lysed in buffer (25 mM HEPES-NaOH, pH 7.4, 150 mM NaCl, 10 mM imidazole, 8.6% glycerol, 1 mM DTT) supplemented with 1 mM MgCl₂, 1 mg/mL lysozyme, a complete EDTA-free protease inhibitor, and benzonase. After gentle shaking at 4 °C for 30 min, lysates were ultrasonicated and centrifuged at 20,000 RPM for 30 min (JA-25.50 rotor, Beckman). The supernatant was incubated with Ni-NTA resin (Qiagen) for 1 h at 4 °C, washed with buffer (25 mM HEPES-NaOH, pH 7.4, 150 mM NaCl, 20 mM imidazole, 8.6% glycerol, 1 mM MgCl₂), and eluted with buffer containing 400 mM imidazole. Eluted fractions were exchanged into lysis buffer (1 mM DTT, 1 mM MgCl₂) and subjected to overnight TEV (MPIB protein production core facility) cleavage at 4 °C. VCP was further purified by size-exclusion chromatography with Superose 6 (Cytiva) in SEC buffer (25 mM HEPES-NaOH, pH 7.4, 150 mM NaCl, 1 mM DTT) and stored at -80°C.

For NPLOC4-UFD1 complex purification, cells expressing 6×His-tagged NPLOC4 and 6×His-thrombin-fused UFD1 were mixed during resuspension in lysis buffer with 1 mg/mL lysozyme, protease inhibitors, and benzonase, then gently shaken at 4 °C for 30 min. After centrifugation, the supernatant was incubated with Ni-NTA resin for 1 h at 4 °C, washed, and eluted as described above. The complex was further purified by size-exclusion chromatography with Superdex 200 (Cytiva) in SEC buffer and stored at -80°C.

ATPase activity was measured using the EnzChek Phosphate Assay Kit (Invitrogen). 500 nM NPLOC4-UFD1 with or without 150 nM VCP were incubated with 20 mM MESG and 1 U/mL purine nucleoside phosphorylase in 1× EnzChek buffer. After a 10 min pre-incubation at 30°C, 2 mM ATP was added, and absorbance at 360 nm was recorded every 20 s for 30 min using a Spark plate reader (Tecan). ATPase activity was calculated from the linear absorbance slope, normalized to VCP-UN, and error bars represent s.d. from three replicates.

### Proteasome degradation assay

Htt64Q were induced with muristerone A as described two large cell culture plates and lysed on ice in buffer (0.5ml 1x PBS +1% Triton) containing protease inhibitor cocktail and benzonase. The lysates were passed through a 23-gauge needle to enhance lysis. The lysate was then centrifuged at 1000g at 4°C for 10 min to remove large cellular clumps, and the more soluble 64Q fraction was collected by pushing the lysate through a pre-equilibrated 0.2 μm syringe filter (Millipore). For the 64Q-GFP pull-down, the fraction was incubated with GFP-Trap at 4°C for 4 h followed by washes in the same buffer without protease inhibitors.

The degradation assay was performed on the pull-down in reaction buffer (25mM Tris pH7.4, 10mM MgCl2, 10% glycerol) with freshly added ATP (4 mM) and DTT (1 mM) at 37°C. The purified 20S and 26S proteasomes (ENZO) were activity-certified. The 26S proteasome preparation contained 19S and 20S complexes at a 6:1 ratio, as assessed by Coomassie staining and quantitative mass spectrometry, and was therefore used as 19S. The 20S proteasome, 19S proteasome, and VCP/NPLOC/UFD1 complexes were added at 2 μg. After 0, 2 or 4 h time points, the reactions were aliquoted and stopped by boiling at 95°C in sample buffer for immunoblotting.

### VCP-20S binding assay with peptide competition

Binding assays were performed using the Dynabeads Protein G immunoprecipitation kit (Invitrogen) per the manufacturer’s protocol. VCP antibody (0.5 μg) was conjugated to 50 µl of beads and incubated with 4 µg of purified VCP. For peptide binding, 5 µg of the 20S proteasome was incubated with HbYX peptides (0, 10 µM, 1 mM) in binding buffer (20 mM Tris-HCl pH 7.4, 150 mM NaCl, 2 mM EDTA) with protease inhibitors for 30 min at room temperature The inclusion of epoxomicin, to irreversibly inhibit the 20S, was also tested it did not result in a significant change in the pattern of the results. The mixture was incubated with beads at 4°C for 2 h with gentle rotation, washed, and eluted in sample buffer for immunoblotting.

### Alphafold3 and sequence analyses

Six copies of the human VCP sequences from the UniProt entry P55072 were analyzed with and without ADP or ATP to capture the different states using AlphaFold3. The results agree with published PDB structures. The PSMA subunits of the human 20S proteasome from UniProt (entries: P25786, P25787, P25788, P25789, P28066, P60900, P14818) were analyzed with 1, 3, 6, or 7 copies of the VCP C-terminal peptide (DNDDDLYG). All analyses were performed using the AlphaFold3 online server (https://alphafoldserver.com/).

VCP C-terminal amino acid sequences were downloaded from Uniprot (entries: P55072 *H. sapiens*, Q3ZBT1 *B. taurus*, P23787 *X. laevis*, Q01853 *M. musculus*, Q7KN62 *D. melanogaster*, O05209 *T. acidophilum*, P25694 *S. cerevisiae*) and multiple sequence alignment performed in Clustal Omega (https://www.ebi.ac.uk/jdispatcher/msa/clustalo).

### Quantification and Statistical analysis

To present quantitative data, Microsoft Excel v.16.0.10406.20006 and GraphPad Prism v.9.5.1 were utilized and figures are structured in Adobe Illustrator 2023. Western blot data displayed were representative experiments, and the number of independent experiments from biological replicates conducted is indicated in the legend.

### Key resources table

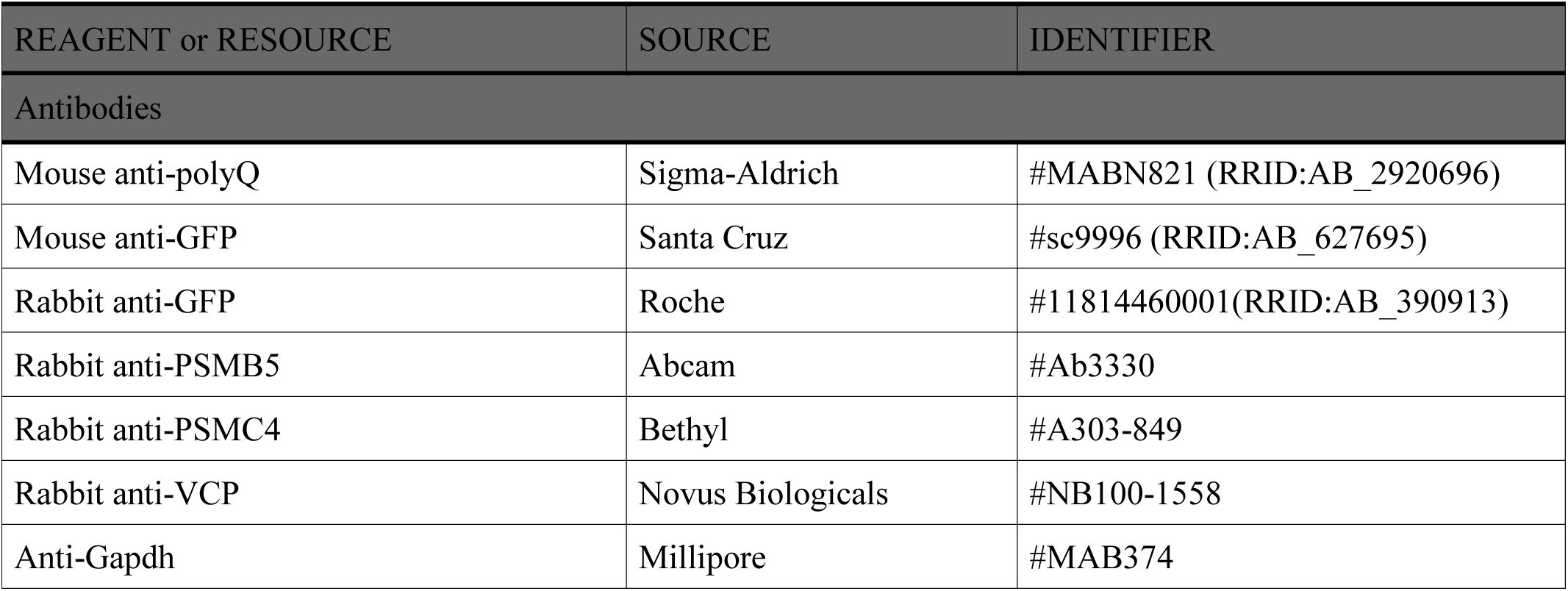
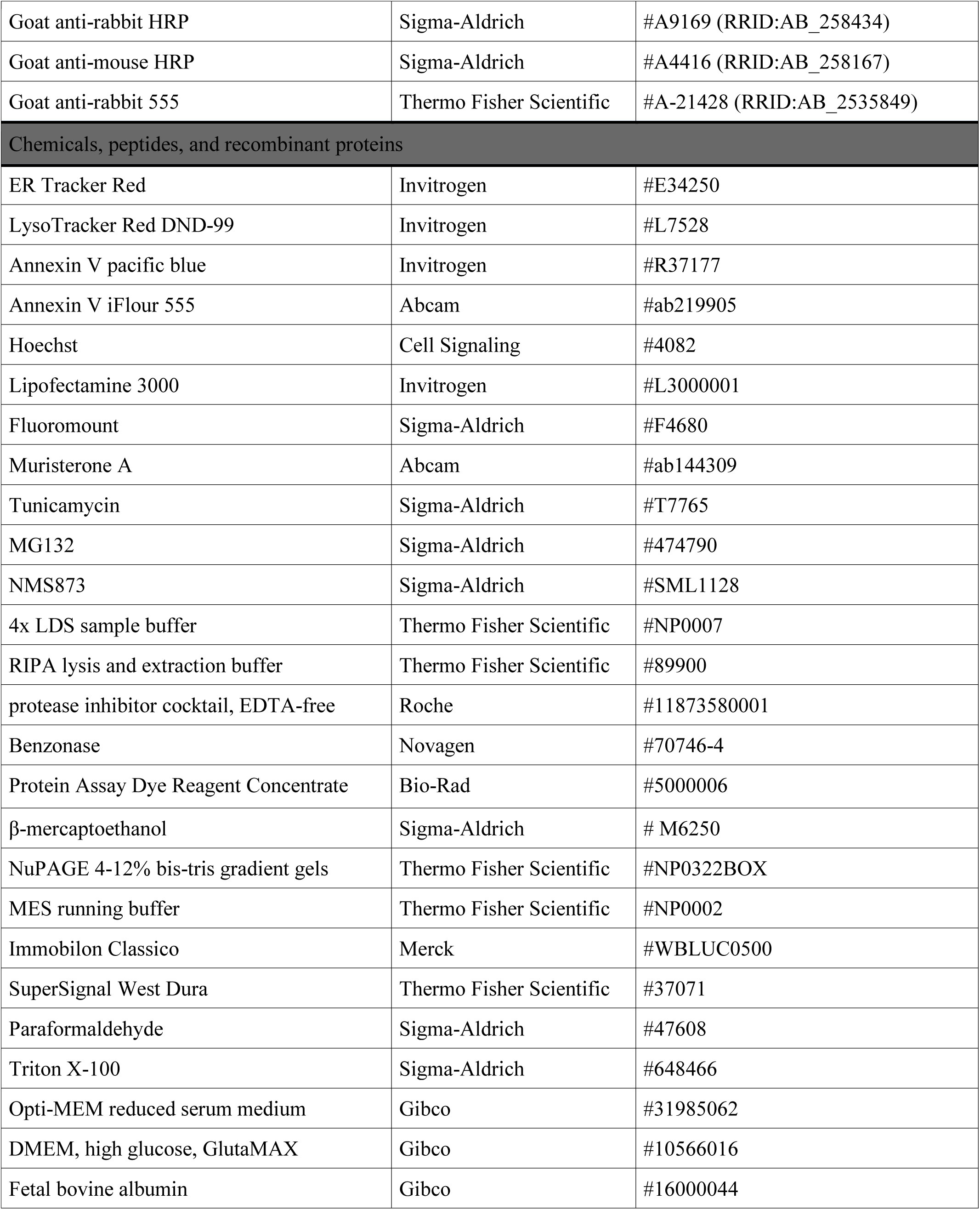
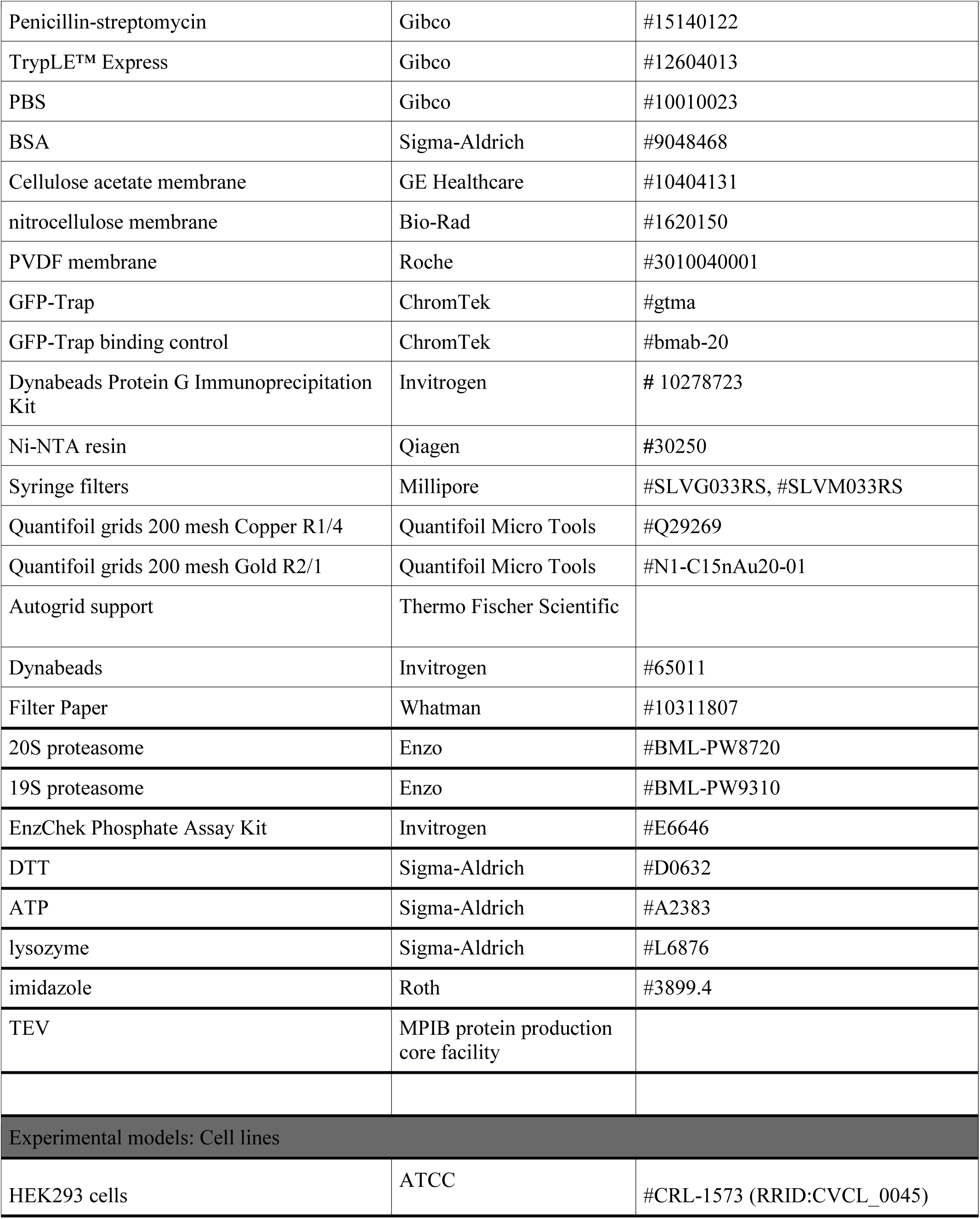
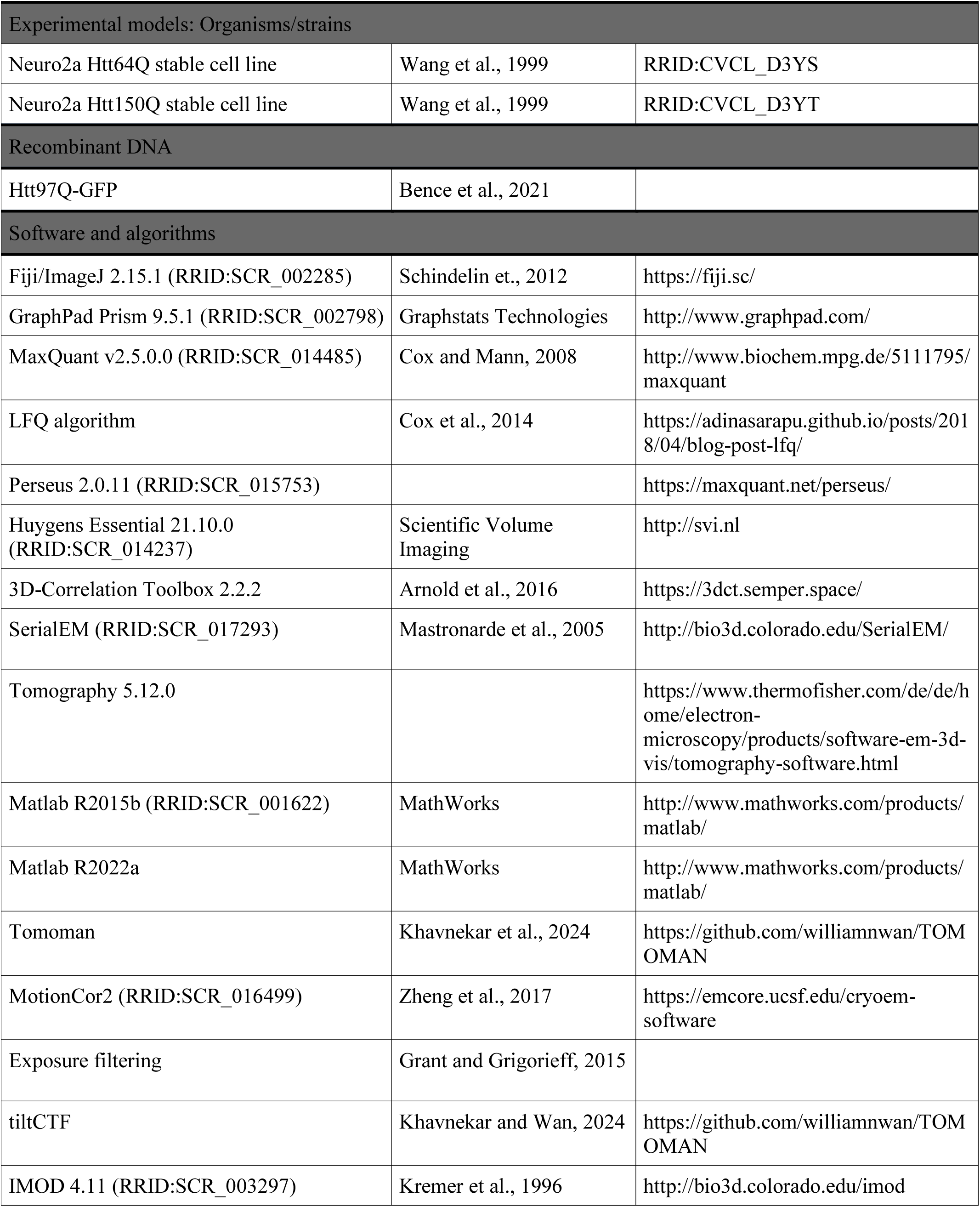
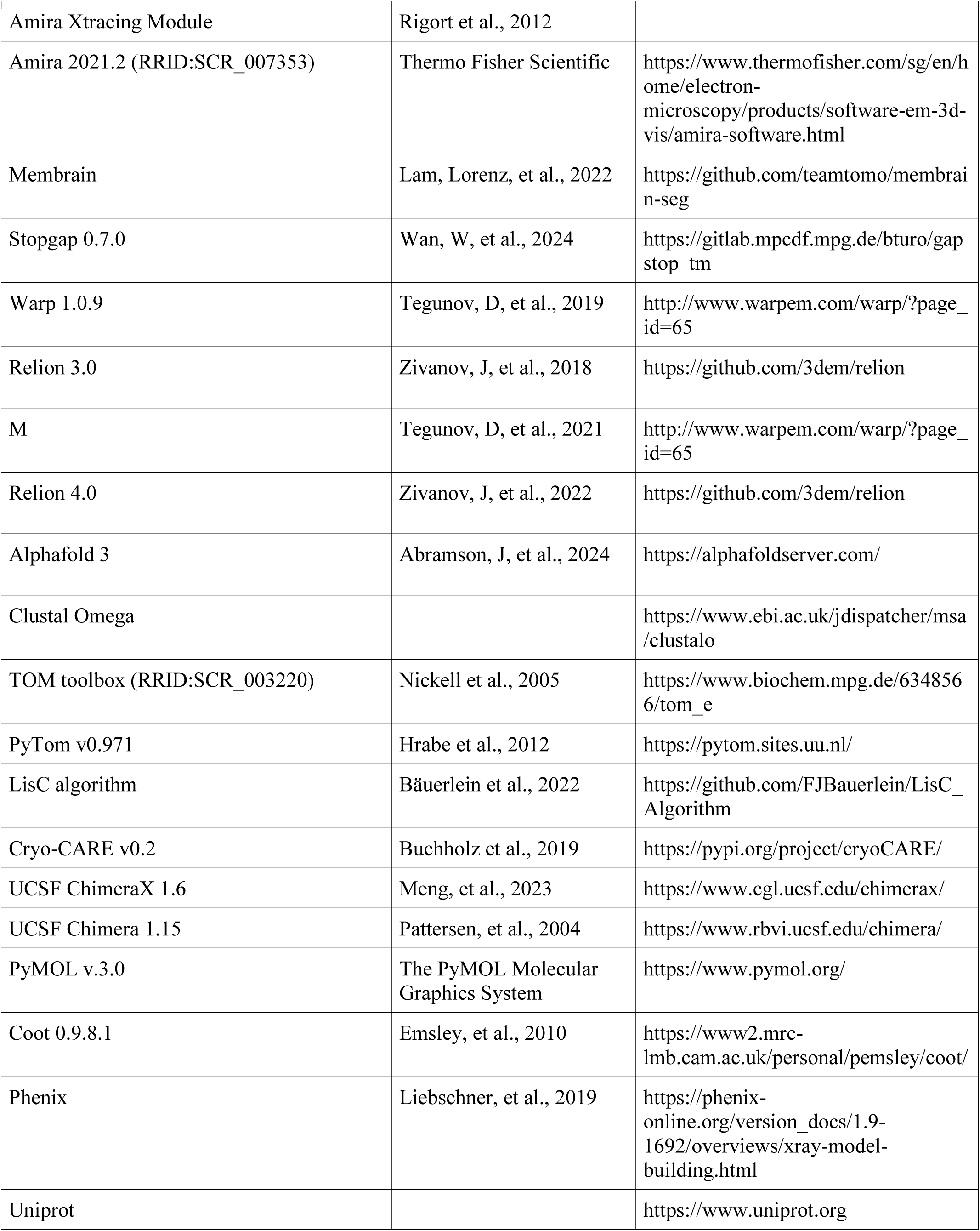
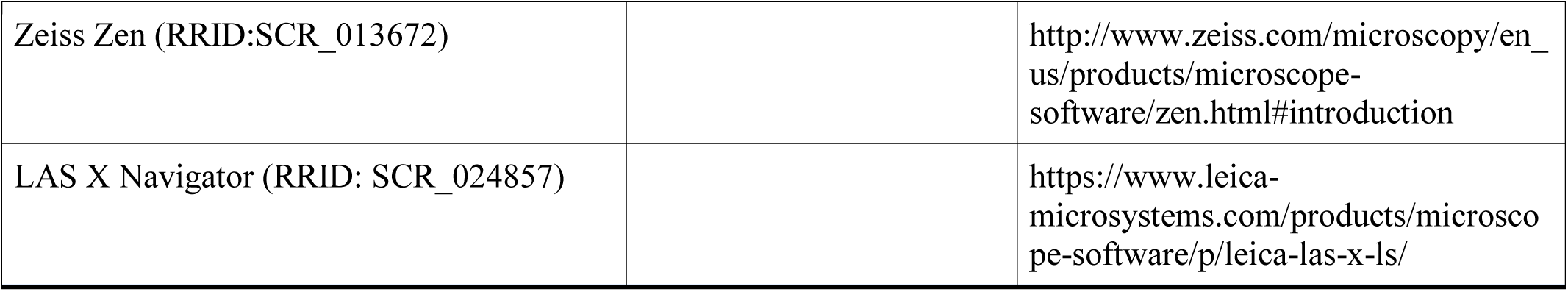

## Acknowledgements

D.Y.Z, and W.B designed the study. W.B. and J.P provided the instruments. D.Y.Z. carried out computational analysis with input from J.W., F.B., C.M., W.J., Q.G., M.P., and P.X.. C.M. purified the VCP and cofactors. D.Y.Z carried out experiments with input from C.M. and I.S.. W.J. wrote the scripts for the nearest distance analysis and distance simulation between the macromolecules, P.X. wrote the script for the nearest distance analysis between macromolecules and fibrils. D.Y.Z. wrote the manuscript with input from everyone. We acknowledge Dr. Franz-Ulrich Hartl at MPI, Tillman Schäfer and Daniel Bollschweiler at MPI EM facility, Martin Spitaler and Markus Oster at the MPI Imaging facility, Barbara Steigenberger and Nicole Krombholz at the MPI mass spectrometry facility, Stephan Uebel and Stephan Pettera at the MPI biochemistry core for input. We acknowledge Hou Zhen, Dustin Morado, Zunlong Ke, Hui Guo, Oda Schiøtz, Sven Klumpe, and Inga Wolf for technical support.

