## Supplementary figures 1-10 for "The targeting of non-fibrillar polyQ via distinct VCP-proteasome coupling"

A

B

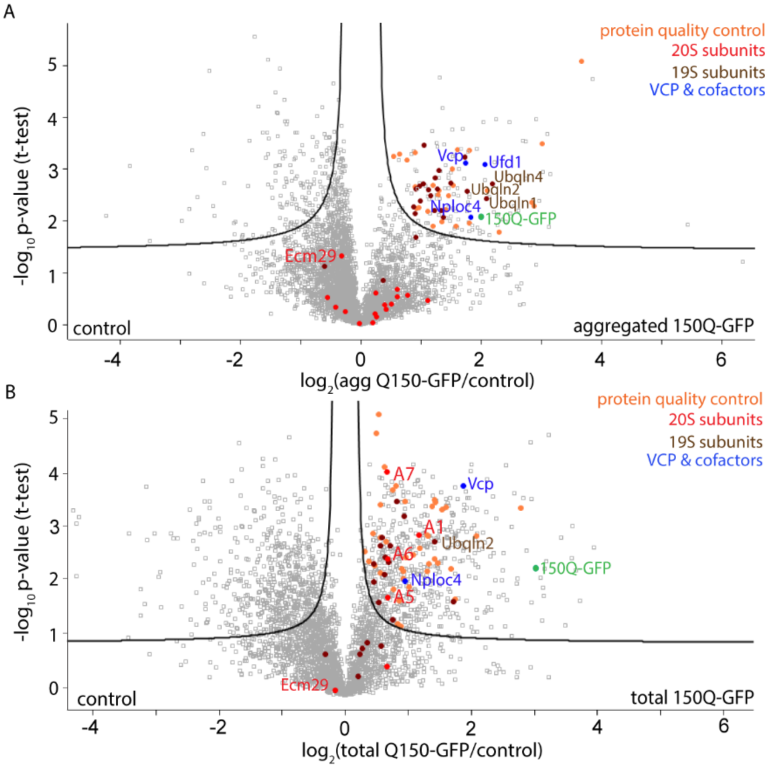


C

| Agg 150Q-GFP | | Total 150Q-GFP | |
|---|---|---|---|
| 19S | 20S | 19S | 20S |
| Psmc1 | | Psmc1 | Psma1 |
| Psmc2 | | Psmc2 | Psma5 |
| Psmc3 | | Psmc4 | Psma6 |
| Psmc4 | | Psmc5 | Psma7 |
| Psmc5 | | Psmd2 | |
| Psmc6 | | Psmd3 | |
| Psmd1 | | Psmd4 | |
| Psmd2 | | Psmd7 | |
| Psmd3 | | Psmd8 | |
| Psmd4 | | Psmd11 | |
| Psmd5 | | Psmd13 | |
| Psmd7 | | Psmd14 | |
| Psmd8 | | | |
| Psmd11 | | | |
| Psmd12 | | | |
| Psmd13 | | | |
| Psmd14 | | | |

D E F G H I J K

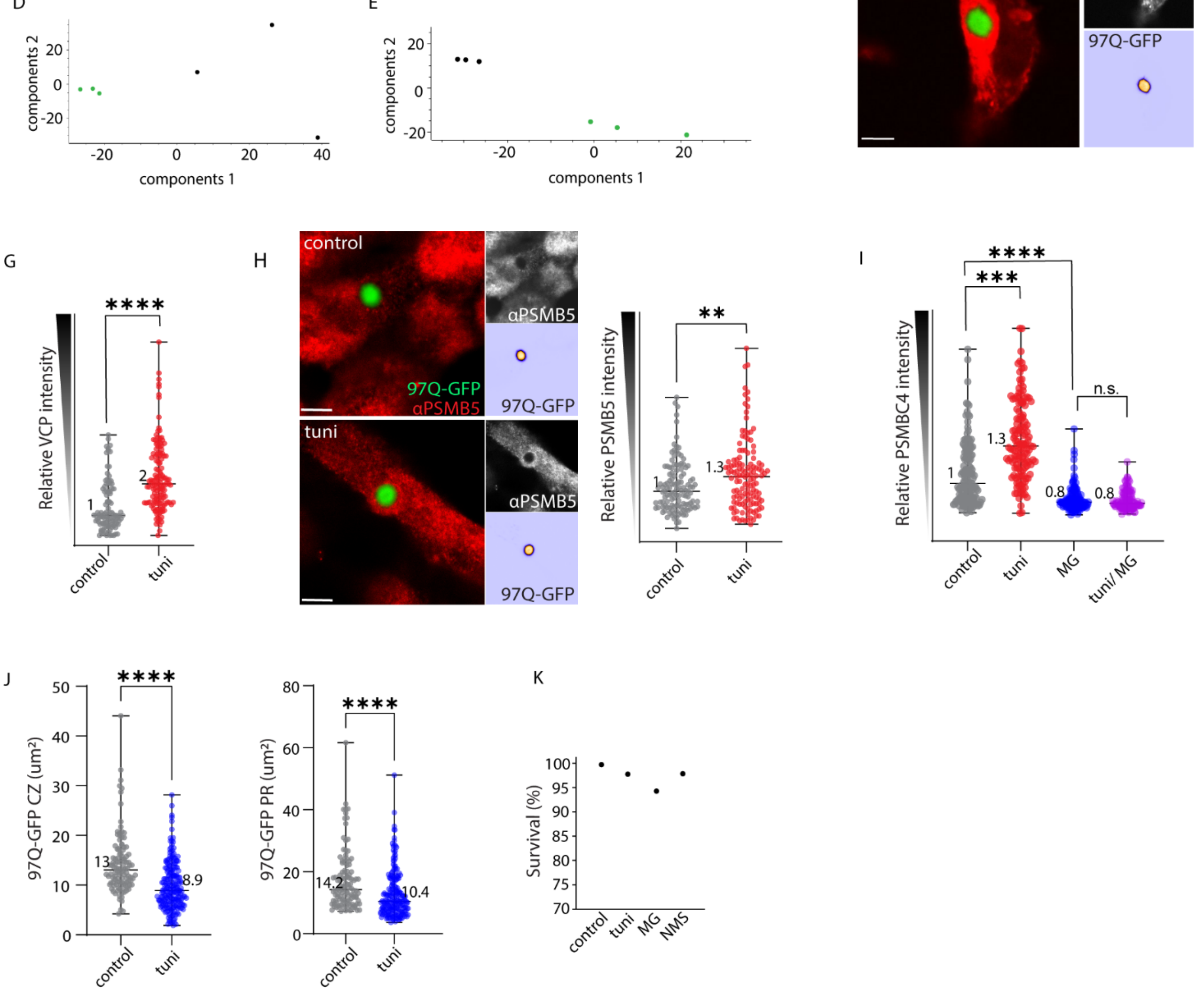

**Figure S1 Non-fibrillar polyQ intermediates are targeted by VCP and proteasome**

**(A-E)** Label-free quantitative mass spectrometry analyses of GFP pull-down from filter-partitioned (> 0.2 μm) 150Q-GFP aggregates (**A**) or total lysate (**B**) from Neuro2a cells. Negative controls were pull-downs using control beads. Curves represent FDR < 0.05. 20S subunits and Ecm29 are highlighted in red, 19S subunits in brown, VCP and cofactors in blue, and other protein quality control factors in orange.
**(C)** A list of the detected 19S and 20S subunits enriched with filter-partitioned aggregated 150Q or total 150Q following GFP pull-down. In the aggregated fraction, 17/20 19S subunits were enriched, while 0/14 20S subunits showed enrichment. (**D-E**) Quantitative mass spectrometry principal component analyses of filter-partitioned fraction (**D**) or from total lysate (**E**) with GFP (green) or control (black) pull-downs in triplicates.

**(F)** Representative confocal image of 97Q-GFP expression in HEK293, co-stained with ER-Tracker.
**(G)** Image quantification of VCP relative intensity within 1.5 μm around the 97Q central zone (CZ) in control or following 9 h induction of ER stress with tuni (2 μg/ml) (control: n = 102, tuni: n = 139).
**(H)** Representative confocal images of 97Q control or treatment in HEK293, co-stained with PSMB5 antibody. Image quantification of PSMB5 relative intensity within 1.5 μm around the 97Q CZ in control and 9 h treatment (control: n = 122, tuni: n = 130).
**(I)** Image quantification of PSMC4 relative intensity within 1.5 μm around the 97Q CZ in control and 9 h treatments (control: n = 141, tuni: n = 137, MG132: n = 130, tuni/MG132: n = 129).
Median values are displayed (**G-I**), intensity standardized with threshold 800-8000 for PR, and > 8000 for CZ, ****p < 0.0001, *** p < 0.001, ** p < 0.01, p values generated from two-tailed student's t-test.
**(J)** Image quantification of 97Q-GFP in HEK293 LC3A/B KO cells, in the CZ and peripheral region (PR) in control and 9 h treatment (CZ: control: n = 111, tuni: n = 104; PR: control: n = 111, tuni: n = 104).
**(K)** Fluorescence live imaging for cell viability in HEK293 upon Annexin V-iFlor 555 staining, control: n = 340, tuni: n = 505, MG132: n = 327, and NMS873: n = 265.
Scale bar: 5 μm in (**F, H**).

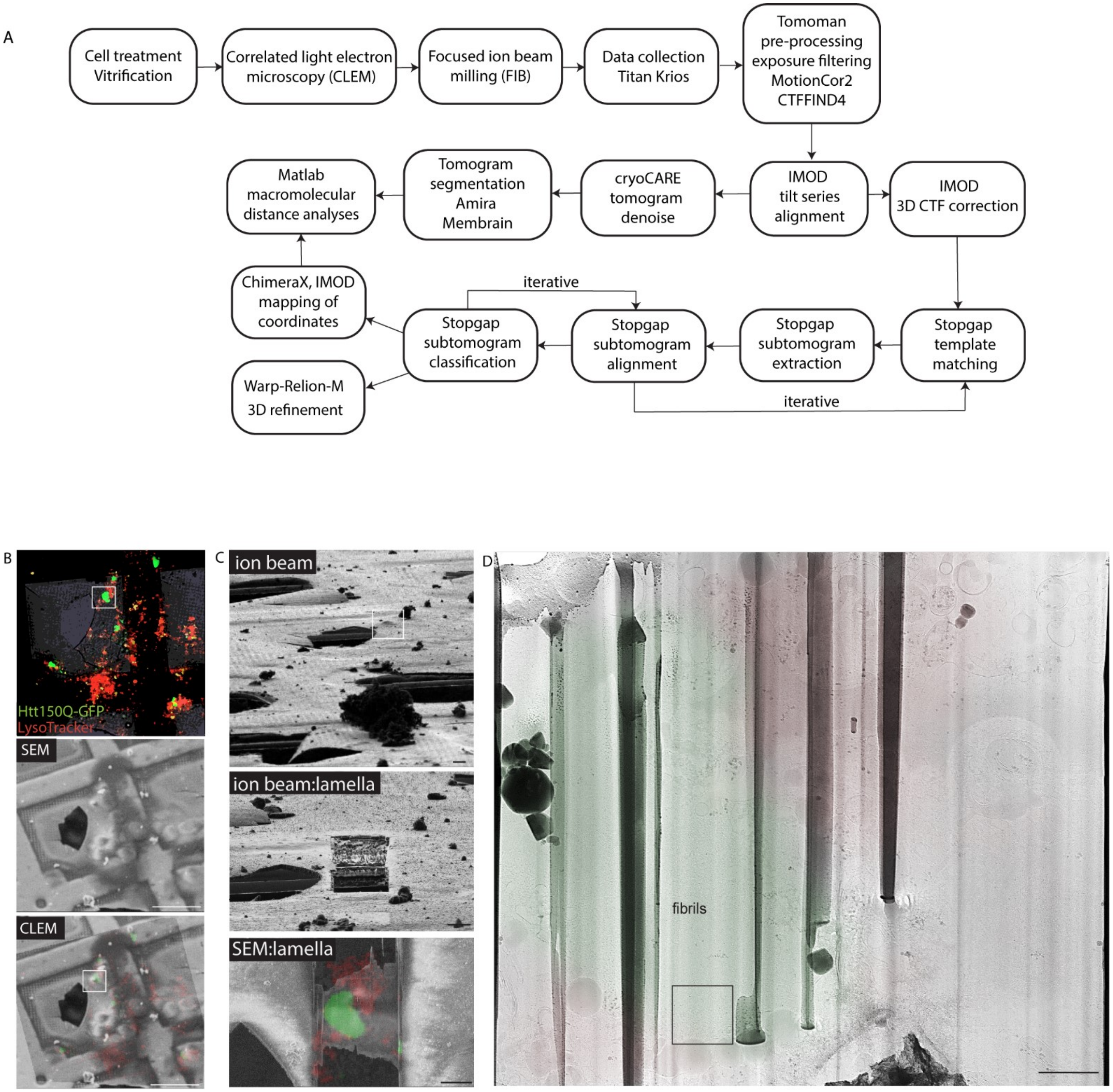

A
Cell treatment
Vitrification
Correlated light electron
microscopy (CLEM)
Focused ion beam
milling (FIB)
Data collection
Titan Krios
Tomoman
pre-processing
exposure filtering
MotionCor2
CTFFIND4
Matlab
macromolecular
distance analyses
Tomogram
segmentation
Amira
Membrain
cryoCARE
tomogram
denoise
IMOD
tilt series
alignment
IMOD
3D CTF correction
iterative
ChimeraX, IMOD
mapping of
coordinates
Stopgap
subtomogram
classification
Stopgap
subtomogram
alignment
Stopgap
subtomogram
extraction
Stopgap
template
matching
Warp-Relion-M
3D refinement
iterative
B
Htt150Q-GFP
LysoTracker
SEM
CLEM
C
ion beam
ion beam:lamella
SEM:lamella
D
fibrils

**Figure S2 *In situ* visualization of VCP and proteasomes around polyQ aggregates**

(**A**) Cryo-CLEM sample preparation, data collection and processing workflow, including the software used for each step.

Related to Figure 2 (dataset 1)
(**B**) Cryo-CLEM workflow: 150Q expressed in Neuro2a stained with LysoTracker (to locate the cytosolic aggregates), site of interest is boxed.
(**C**) FIB and SEM views of the target, site of interest is boxed.
(**D**) TEM Lamella view (8700x) showing the tomogram area (boxed) overlaid with fluorescence signal.
Scale bars: 50 μm (**B**), 5 μm (**C**), 1 μm (**D**).

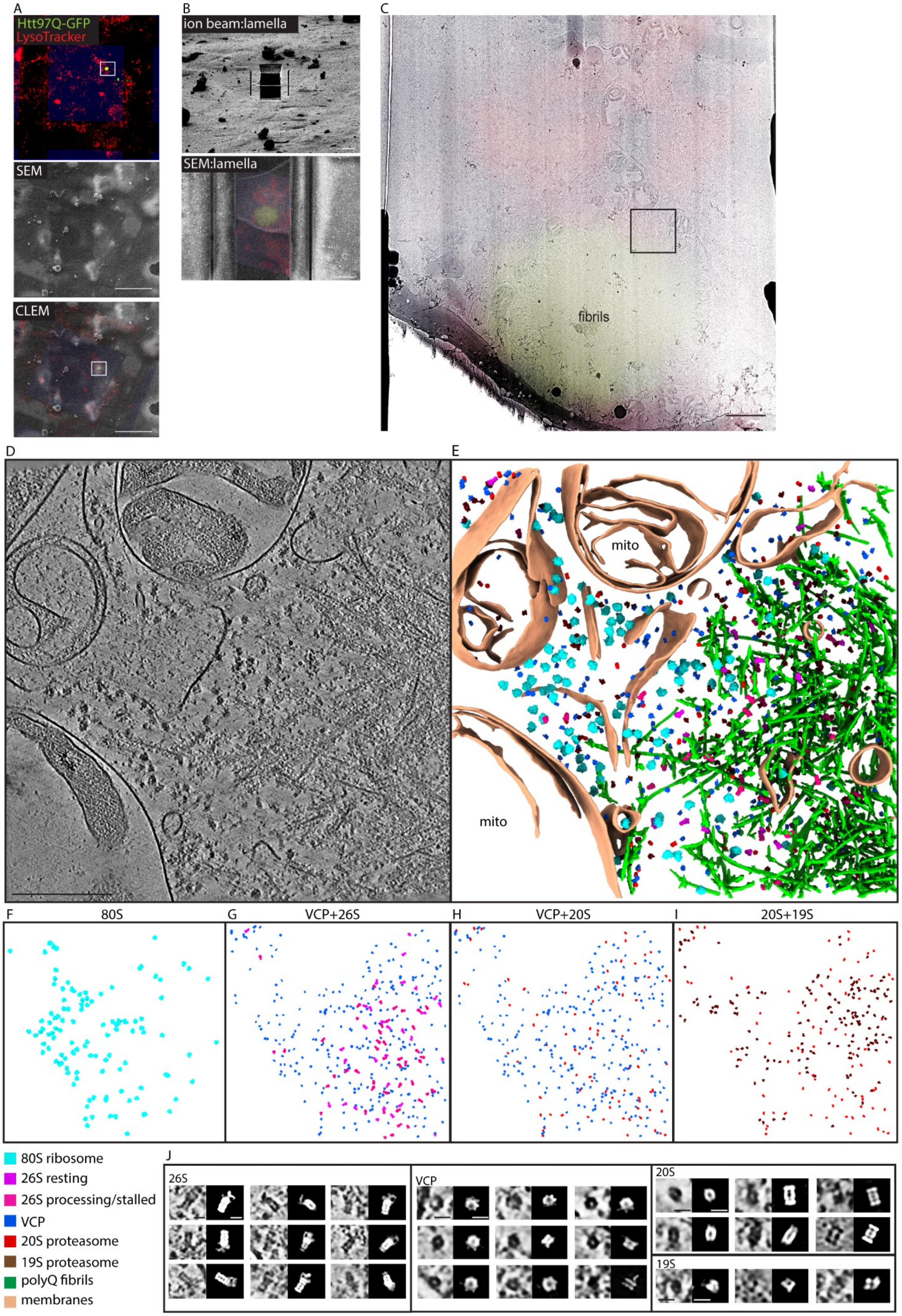

A
Htt97Q-GFP
LysoTracker
SEM
CLEM
B
ion beam:lamella
SEM:lamella
C
fibrils
D
E
mito
mito
F
80S
G
VCP+26S
H
VCP+20S
I
20S+19S
80S ribosome
26S resting
26S processing/stalled
VCP
20S proteasome
19S proteasome
polyQ fibrils
membranes
J
26S
VCP
20S
19S

**Figure S3 *In situ* visualization of VCP and proteasomes around polyQ aggregates**

Related to Figure 2 (dataset 1)
(**A**) Cryo-CLEM workflow: 97Q expressed in HEK293 stained with LysoTracker, site of interest is boxed.
(**B**) FIB and SEM views of the target.
(**C**) TEM Lamella view (8700x) showing the tomogram area (boxed) overlaid with fluorescence signal.
**(D-I)** A slice of denoised tomogram (**D**) and the 3D renderings (**E-I**) illustrating macromolecular distributions. Mito: mitochondria.
(**J**) Enlarged views of the macromolecules detected in the tomogram (**D**) with their coordinates determined by Stopgap.
Scale bars: 50 µm (**A**), 5 µm (**B**), 1 µm (**C**), 250 nm (**D**), 15 nm (**J**).

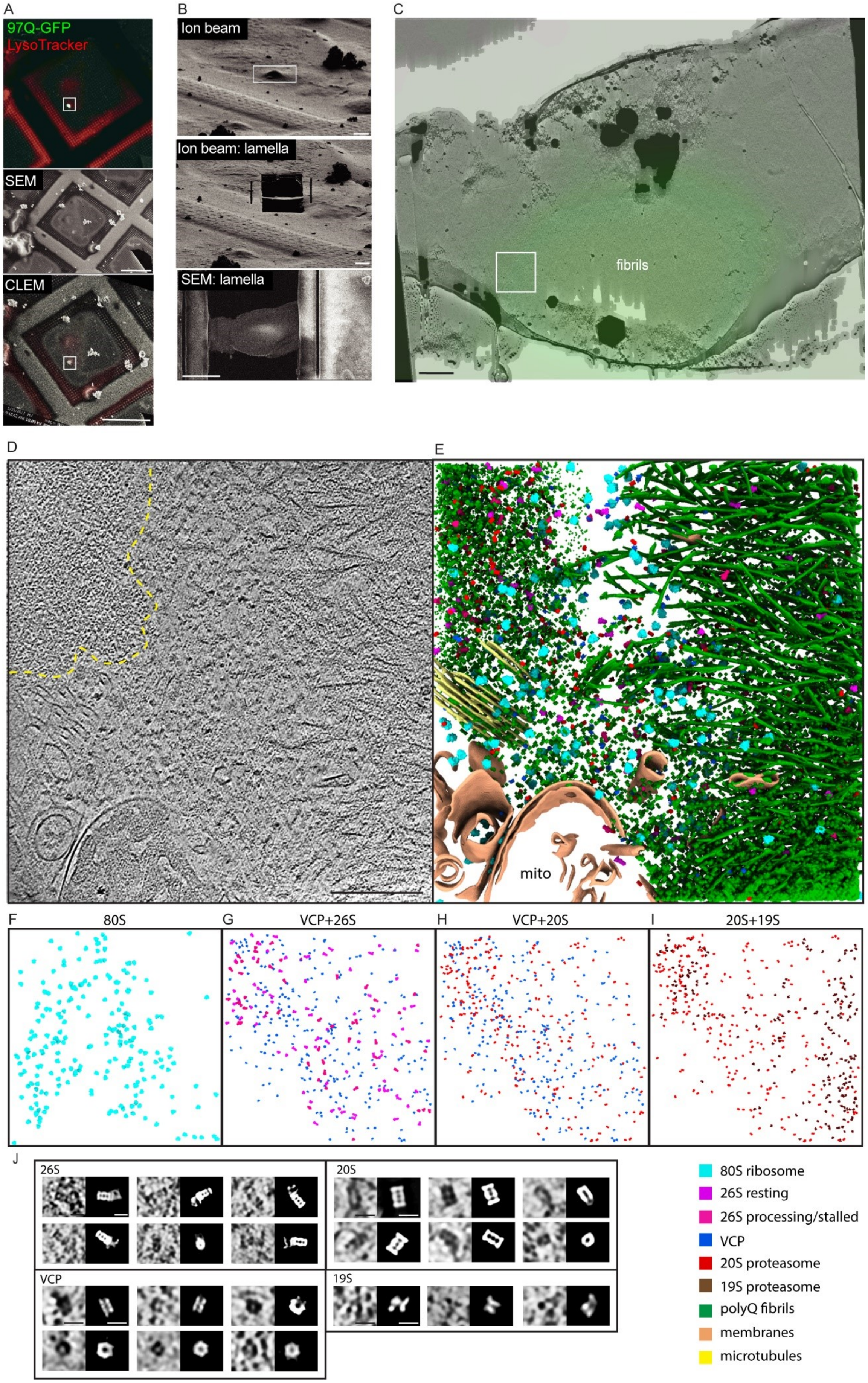
A
97Q-GFP
LysoTracker
SEM
CLEM
B
Ion beam
Ion beam: lamella
SEM: lamella
C
fibrils
D
E
mito
F
80S
G
VCP+26S
H
VCP+20S
I
20S+19S
J
26S
20S
VCP
19S
80S ribosome
26S resting
26S processing/stalled
VCP
20S proteasome
19S proteasome
polyQ fibrils
membranes
microtubules

**Figure S4 *In situ* visualization of VCP and proteasomes around polyQ aggregates**

Related to Figure 2 (dataset 1)
(**A**) Cryo-CLEM workflow: 97Q expressed in HEK293 LC3A/B KO stained with LysoTracker, site of interest is boxed.
(**B**) FIB and SEM views of the target, site of interest boxed.
(**C**) TEM Lamella view (5600x) showing the tomogram area (boxed) overlaid with fluorescence signal.
**(D-I)** A slice of denoised tomogram (**D**) and 3D renderings (**E-I**) illustrating macromolecular distributions, fibrillar and amorphous polyQ are segmented in green (**E**). The dense amorphous polyQ region is indicated by dotted line in (**D)**.
(**J**) Enlarged views of the macromolecules detected in the tomogram (**D**) with their coordinates determined by Stopgap.
Scale bars: 50 µm (**A**), 5 µm (**B**), 1 µm (**C**), 250 nm (**D**), 15 nm (**J**).

A

| Dataset 1: 250 tomograms | 80S ribosome | 26S resting | 26S processing | 26S state uncertain | 20S | 19S | VCP (ADP) | VCP (ATP) | VCP-cofactor |
|---|---|---|---|---|---|---|---|---|---|
| Microscope/Voltage | Titan Krios 300kV | | | | | | | | |
| Camera | Gatan K2 Summit | | | | | | | | |
| Energy filter | 20 eV | | | | | | | | |
| Software | SerialEM | | | | | | | | |
| tilt angle and tilt span | 2°, ~120 | | | | | | | | |
| Exposure (electrons/Å² ) | 80-120 | | | | | | | | |
| Defocus range | 3-5 μm | | | | | | | | |
| Pixel size (Å) | 3.35 | | | | | | | | |
| Symmetry (Å) | C1 | C1 | C1 | C1 | D7 | C1 | C6 | C1 | C1 |
| Subtomograms (initial extract) | 114389 | 98982 | | | 107264 | 68901 | 117125 | 90879 | 60148 |
| Subtomograms (final) | 83309 | 7562 | 4223 | 9374 | 29314 | 25588 | 8331 | 23749 | 23091 |
| Intial model (PDB) | 4UG0 | 5MPB | | | 6RGQ | 8AMZ | 5FTK | 5FTN | 6CHS |
| FSC threshold | 0.143 | | | | | | | | |
| Relion bin2 resolution (Å) | 13.7 | 20 | 19 | >20 | 15 | 19 | 18 | 17 | 16 |

B

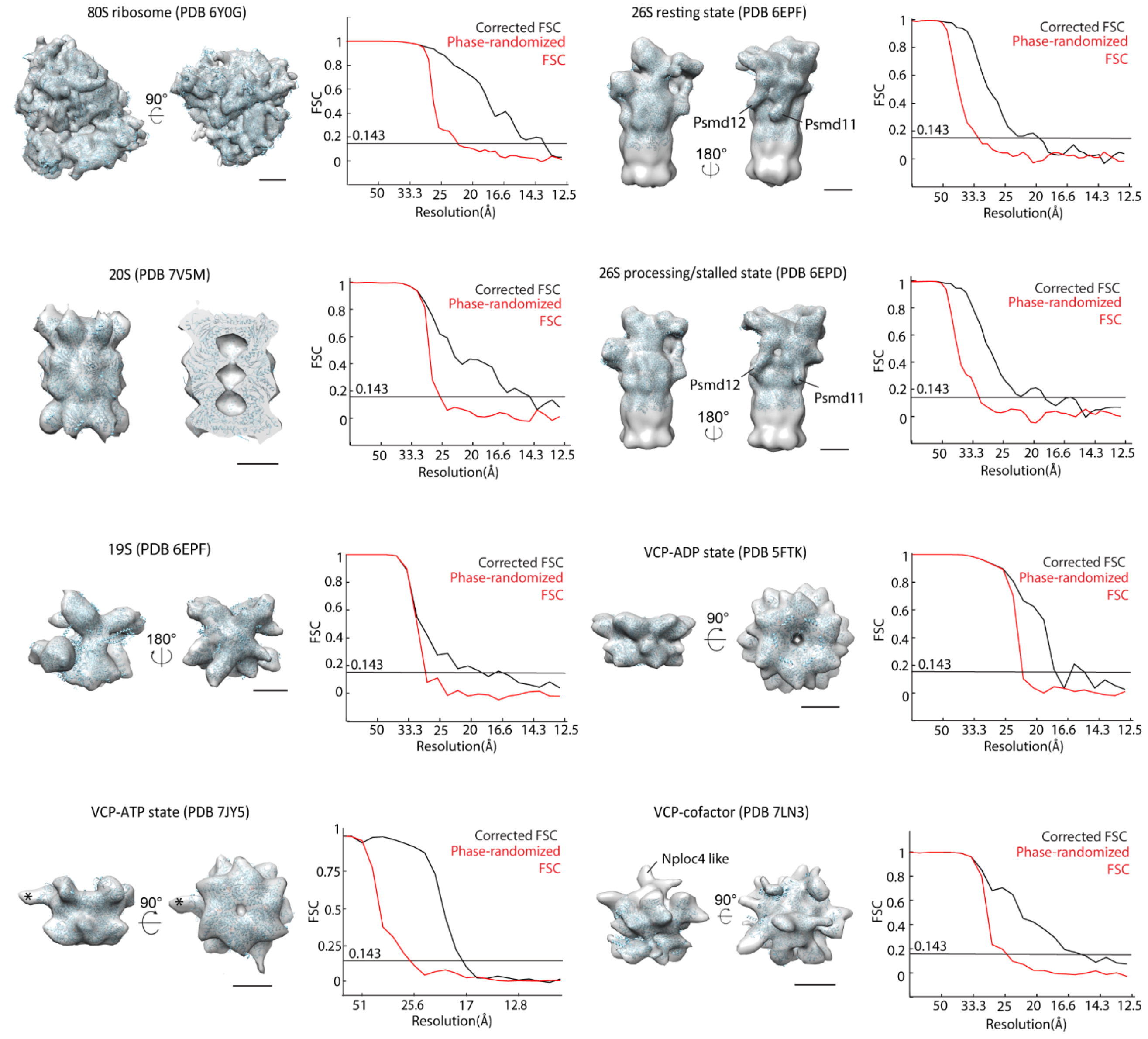

**Figure S5 *In situ* cryo-ET subtomogram averaging (dataset 1)**

(**A**) Datset 1 instrument and data specifications, and resolution estimates.
(**B**) Subtomogram averages of macromolecules, shown as surface representations docked with PDB structures in ribbon view, displayed with corresponding FSC curves. Extra density (*) observed in the VCP-ATP state.
Scale bars: 5 nm (**B**).

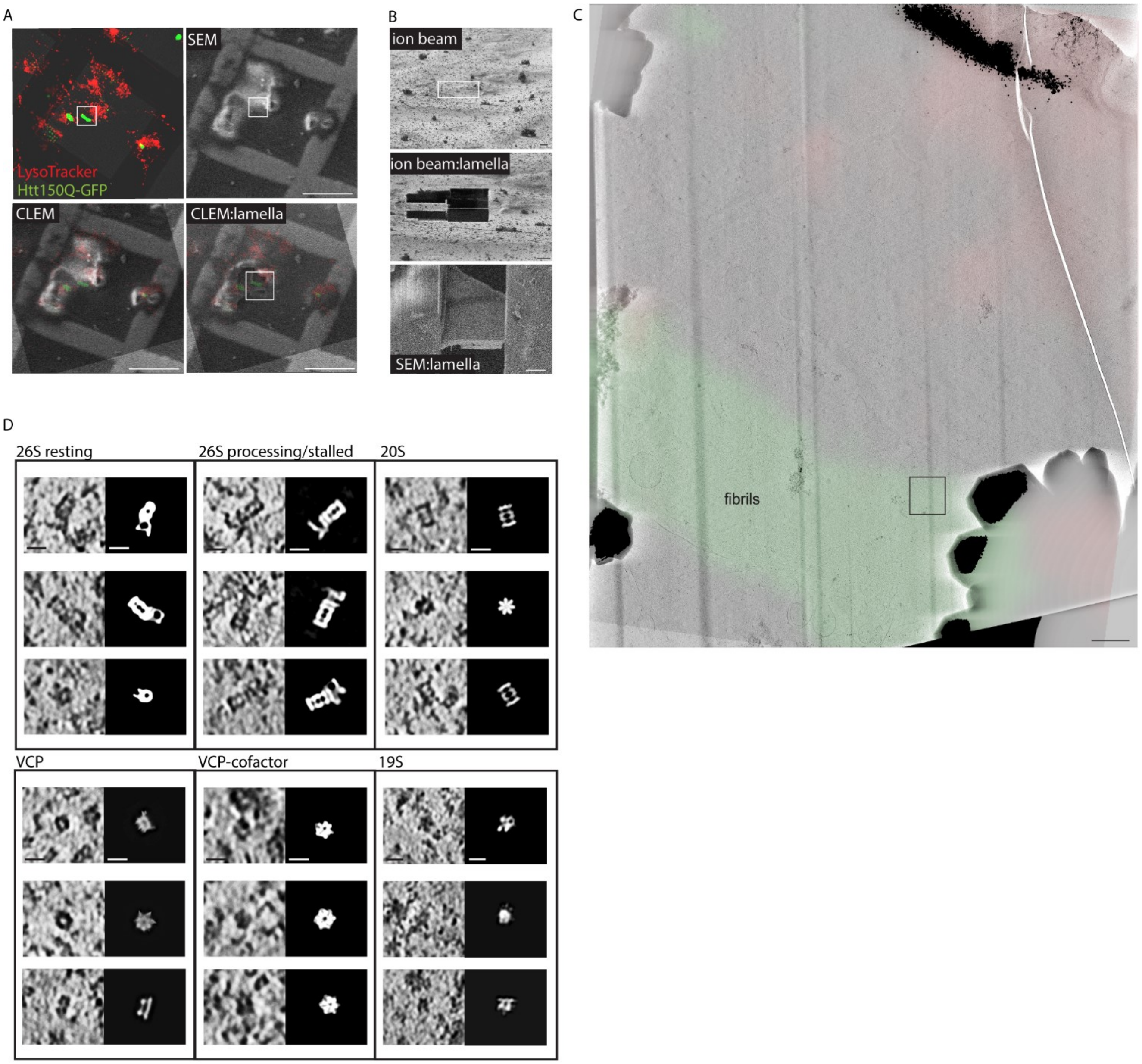


E

| Dataset 2: 410 tomograms | 80S ribosome | 26S resting | 26S processing | 20S | 19S | VCP (ADP) | VCP (ATP) | VCP-cofactor |
|---|---|---|---|---|---|---|---|---|
| Microscope/Voltage | Titan Krios 300kV | | | | | | | |
| Camera | EF-Falcon 4i | | | | | | | |
| Energy filter | 10 eV | | | | | | | |
| Software | Tomography 5 | | | | | | | |
| tilt angle and tilt span | 3°, ~120 | | | | | | | |
| Exposure (electrons/Å$^2$ ) | 80-120 | | | | | | | |
| Defocus range | 3-5 µm | | | | | | | |
| Pixel size (Å) | 1.89 | | | | | | | |
| Symmetry | C1 | C1 | C1 | D7 | C1 | C6 | C1/ C6 | C1 |
| Subtomograms (initial extract) | 46517 | 69373 | | 60503 | 92702 | 74616 | 63921 | 89695 |
| Subtomograms (final) | 45570 | 6443 | 5541 | 16146 | 17476 | 3463 | 19637 | 20719 |
| Intial model (low-pass filtered) | dataset1 reconstruction | hybrid of dataset1 reconstruction | | dataset1 reconstruction | dataset1 reconstruction | hybrid of dataset1 reconstruction | | EMD 23451 |
| FSC threshold | 0.143 | | | | | | | |
| Relion bin2 resolution (Å) | 7.9 | 13 | 11 | 10 | 14 | 14 | 10/9.2 | 11 |

**Figure S6 *In situ* reconstructions with improved resolution for macromolecular sociology**

Related to Figure 3 (dataset 2)
(**A**) Cryo-CLEM: 150Q expressed in Neuro2a stained with LysoTracker, site of interest is boxed.
(**B**) FIB and SEM views of the target, site of interest boxed.
(**C**) TEM Lamella view (11500x) indicating the tomogram area, overlaid with fluorescence signal.
(**D**) Enlarged views of the macromolecules detected in the tomogram (Figure 3A), alongside their coordinates from Stopgap.
Scale bars: 50 µm (**A**), 5 µm (**B**), 1 µm (**C**), 10 nm (**D**).
(**E**) Dataset 2 instrument and data specifications, and resolution estimates.

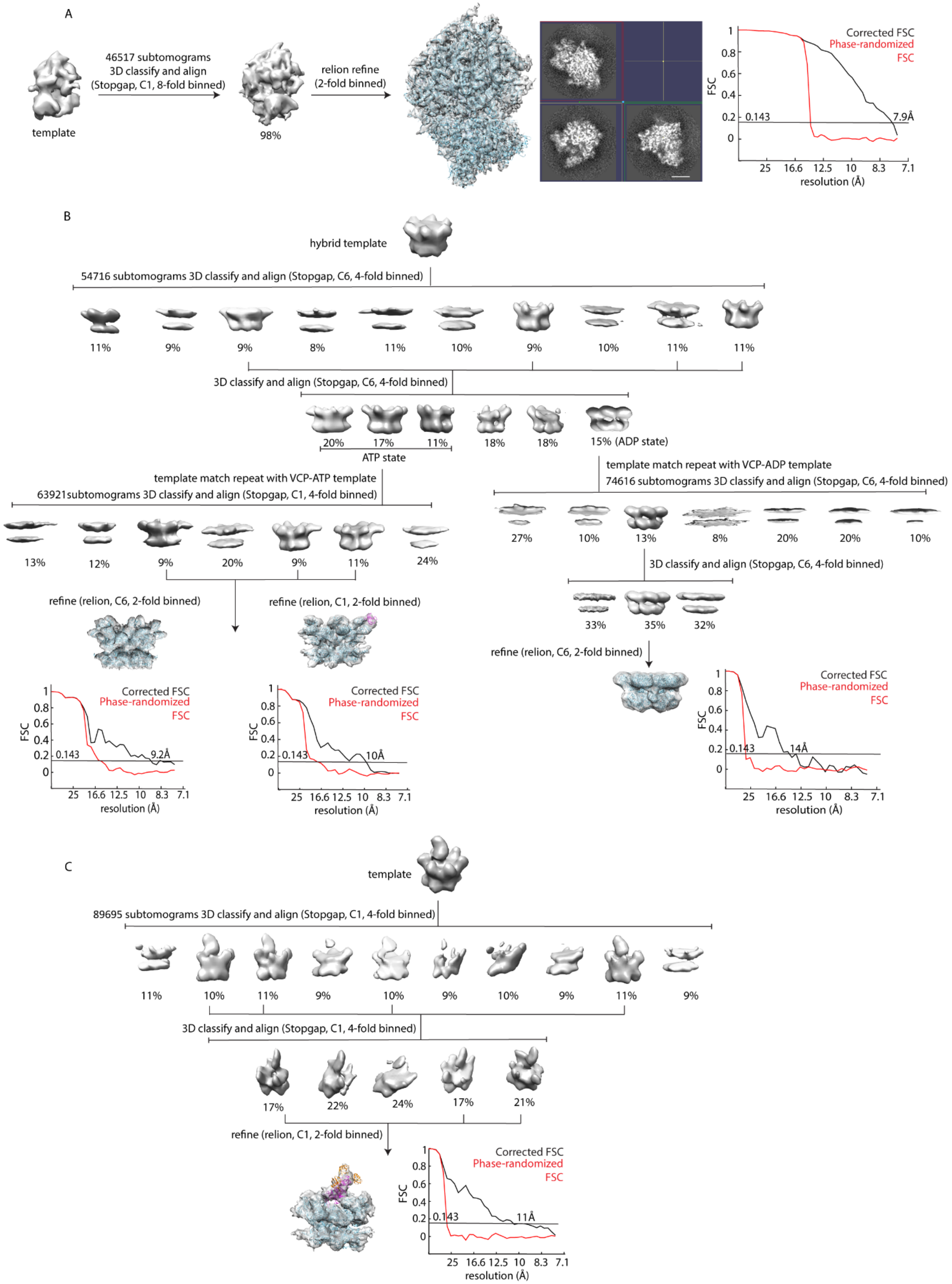
A
template
46517 subtomograms
3D classify and align
(Stopgap, C1, 8-fold binned)
98%
relion refine
(2-fold binned)
Corrected FSC
Phase-randomized FSC
FSC
0.143
7.9Å
resolution (Å)
B
hybrid template
54716 subtomograms 3D classify and align (Stopgap, C6, 4-fold binned)
11% 9% 9% 8% 11% 10% 9% 10% 11% 11%
3D classify and align (Stopgap, C6, 4-fold binned)
20% 17% 11% 18% 18% 15% (ADP state)
ATP state
template match repeat with VCP-ATP template
63921subtomograms 3D classify and align (Stopgap, C1, 4-fold binned)
13% 12% 9% 20% 9% 11% 24%
template match repeat with VCP-ADP template
74616 subtomograms 3D classify and align (Stopgap, C6, 4-fold binned)
27% 10% 13% 8% 20% 20% 10%
3D classify and align (Stopgap, C6, 4-fold binned)
33% 35% 32%
refine (relion, C6, 2-fold binned)
refine (relion, C1, 2-fold binned)
refine (relion, C6, 2-fold binned)
Corrected FSC
Phase-randomized FSC
0.143
9.2Å
10Å
14Å
resolution (Å)
C
template
89695 subtomograms 3D classify and align (Stopgap, C1, 4-fold binned)
11% 10% 11% 9% 10% 9% 10% 9% 11% 9%
3D classify and align (Stopgap, C1, 4-fold binned)
17% 22% 24% 17% 21%
refine (relion, C1, 2-fold binned)
Corrected FSC
Phase-randomized FSC
0.143
11Å
resolution (Å)

**Figure S7 *In situ* reconstructions with improved resolution for the 80S ribosome and VCP**

Related to Figure 3 (dataset 2)
**(A)** *In situ* reconstruction workflow of the 80S ribosomes. The reconstruction is superimposed with PDB 6Y0G in ribbon representation, also shown are the XYZ views and the FSC curve of the reconstruction.
**(B)** *In situ* reconstruction workflow of VCP, corresponding to Figure 3E-F, the reconstructions are displayed next to the corresponding FSC curves.
**(C)** *In situ* reconstruction workflow of the VCP-cofactor complex, corresponding to Figure 3G, the reconstruction is displayed next to the corresponding FSC curve.
Scale bar: 10 nm (**A**).

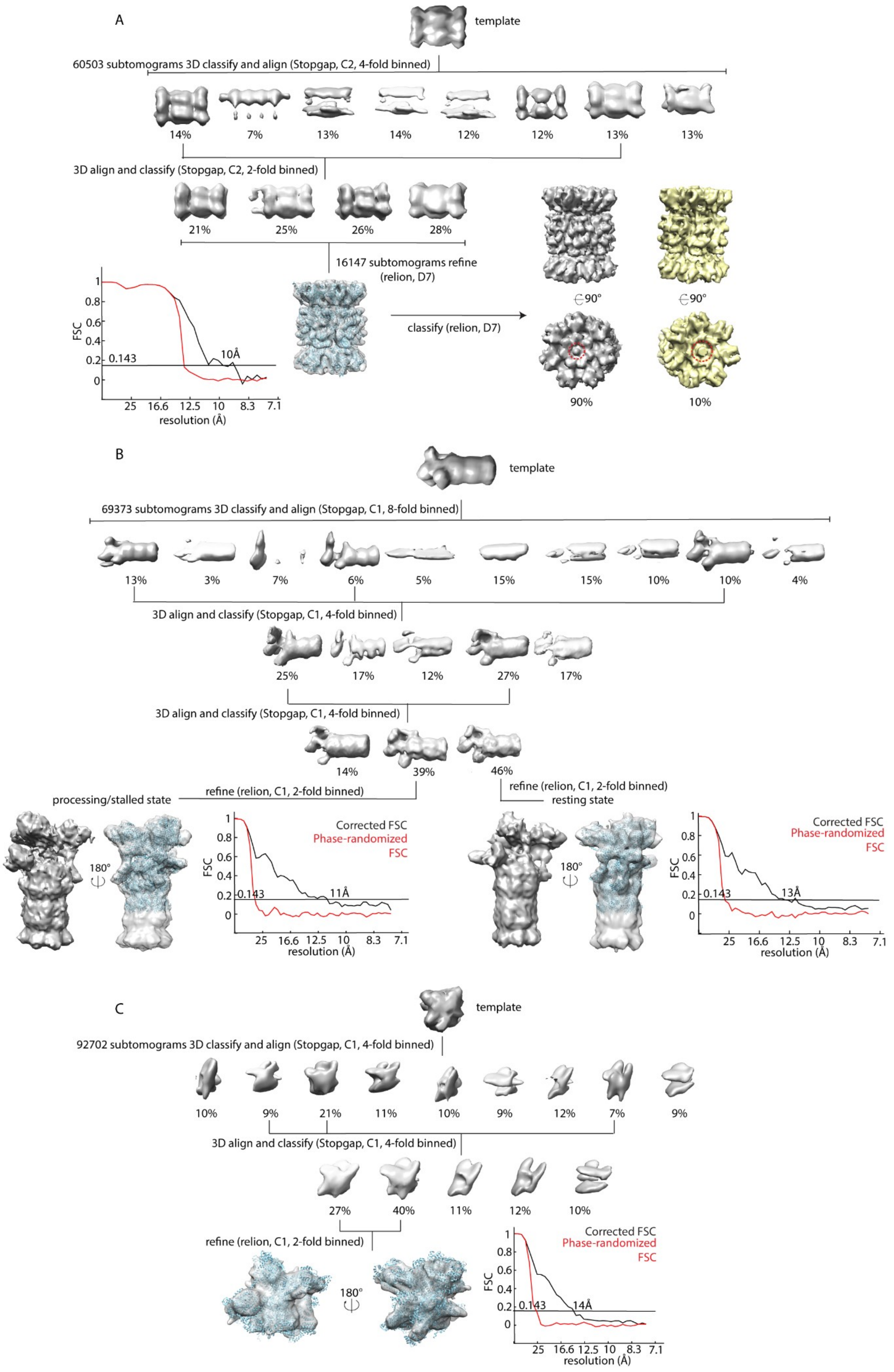

A
template
60503 subtomograms 3D classify and align (Stopgap, C2, 4-fold binned)
14%
7%
13%
14%
12%
12%
13%
13%
3D align and classify (Stopgap, C2, 2-fold binned)
21%
25%
26%
28%
16147 subtomograms refine (relion, D7)
classify (relion, D7)
90°
90°
90%
10%
FSC
1
0.8
0.6
0.4
0.2
0
0.143
10Å
25
16.6
12.5
10
8.3
7.1
resolution (Å)
B
template
69373 subtomograms 3D classify and align (Stopgap, C1, 8-fold binned)
13%
3%
7%
6%
5%
15%
15%
10%
10%
4%
3D align and classify (Stopgap, C1, 4-fold binned)
25%
17%
12%
27%
17%
3D align and classify (Stopgap, C1, 4-fold binned)
14%
39%
46%
refine (relion, C1, 2-fold binned)
processing/stalled state
refine (relion, C1, 2-fold binned)
resting state
180°
Corrected FSC
Phase-randomized FSC
FSC
0.143
11Å
13Å
resolution (Å)
C
template
92702 subtomograms 3D classify and align (Stopgap, C1, 4-fold binned)
10%
9%
21%
11%
10%
9%
12%
7%
9%
3D align and classify (Stopgap, C1, 4-fold binned)
27%
40%
11%
12%
10%
refine (relion, C1, 2-fold binned)
180°
Corrected FSC
Phase-randomized FSC
FSC
0.143
14Å
25
16.6
12.5
10
8.3
7.1
resolution (Å)

**Figure S8 *In situ* reconstructions with improved resolution for the proteasomes**

Related to Figure 3 (dataset 2)
**(A)** *In situ* reconstruction workflow of the 20S proteasome, corresponding to Figure 3H, the reconstruction is shown next to the corresponding FSC curve. Further 3D classification (D7) revealed 10% of the particles to have a different gate configuration, marked by red doted lines.
**(B)** *In situ* reconstruction workflow of the 26S proteasomes, shown with solid surface reconstructions superimposed with PDB structures 6EPF and 6EPD in ribbon representation, along with the corresponding FSC curves.
**(C)** *In situ* reconstruction workflow of the 19S proteasome, shown with solid surface reconstruction superimposed with PDB 6EPF in ribbon representation, along with the corresponding FSC curve.

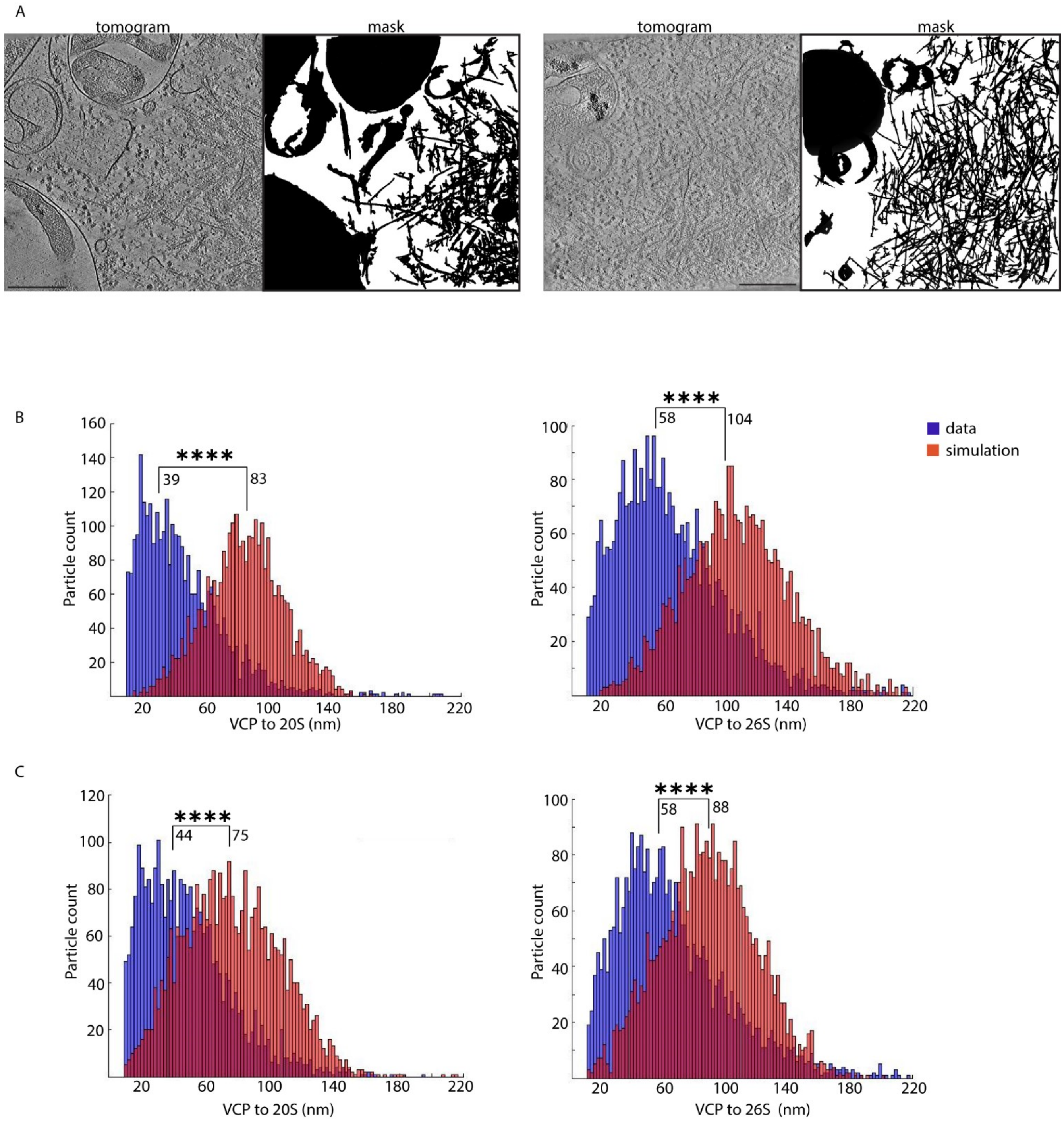
A
tomogram
mask
tomogram
mask
B
****
39
83
Particle count
VCP to 20S (nm)
****
58
104
data
simulation
Particle count
VCP to 26S (nm)
C
****
44
75
Particle count
VCP to 20S (nm)
****
58
88
Particle count
VCP to 26S (nm)

**Figure S9 *In situ* macromolecular sociology suggests cooperativity between VCP and proteasomes**

Analysis related to dataset 1.
**(A)** Examples of masks shown next to the corresponding tomograms, illustrating the removal of regions (black) occupied by fibrils, membranes and organelles (i.e. mitochondria, nucleus). The remaining space (white) is used to distribute macromolecules randomly for the simulation.
(**B, C**) Nearest distance measurements between the particle centers for VCP (regardless of state) and the 20S or 26S proteasomes, compared to random simulation (pink), with a minimum distance cutoff > 10 nm. Median distance values are displayed, ****p < 0.001 Kolmogorov-Smirnov test. Analyses performed on 97Q aggregates in HEK293 (**B**, n = 15 tomograms) and on 150Q aggregates in Neuro2a (**C**, n = 16 tomograms).
Scale bars: 250 nm (**A**).

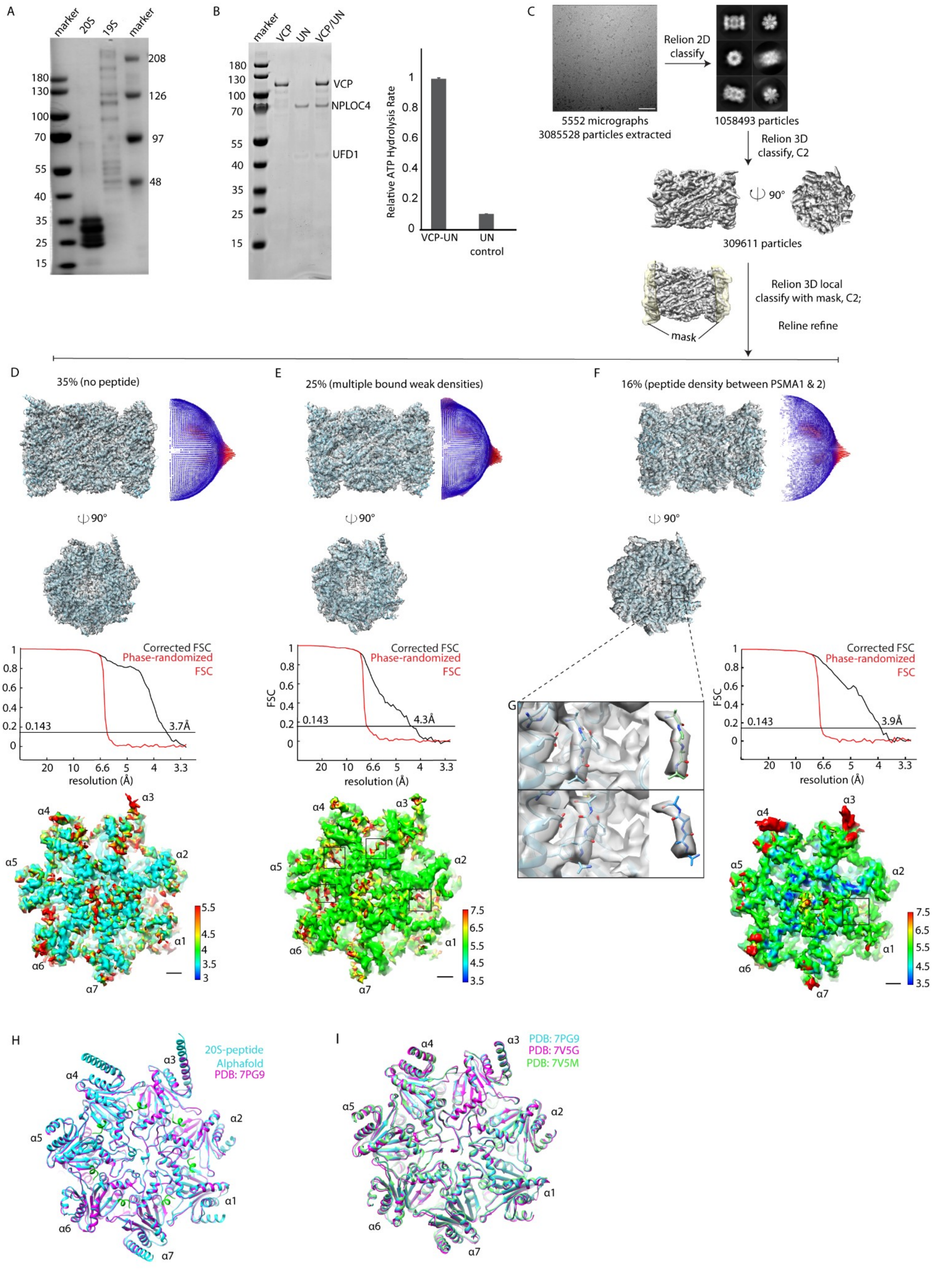

A
marker
20S
19S
marker
180
130
100
70
55
40
35
25
15
208
126
97
48
B
marker
VCP
UN
VCP/UN
180
130
100
70
55
40
35
25
15
VCP
NPLOC4
UFD1
Relative ATP Hydrolysis Rate
1
0.8
0.6
0.4
0.2
0
VCP-UN
UN
control
C
Relion 2D
classify
5552 micrographs
3085528 particles extracted
1058493 particles
Relion 3D
classify, C2
90°
309611 particles
Relion 3D local
classify with mask, C2;
Reline refine
mask
D
35% (no peptide)
90°
Corrected FSC
Phase-randomized
FSC
0.143
3.7Å
resolution (Å)
E
25% (multiple bound weak densities)
90°
Corrected FSC
Phase-randomized
FSC
0.143
4.3Å
resolution (Å)
F
16% (peptide density between PSMA1 & 2)
90°
G
Corrected FSC
Phase-randomized
FSC
0.143
3.9Å
resolution (Å)
α1
α2
α3
α4
α5
α6
α7
H
20S-peptide
Alphafold
PDB: 7PG9
I
PDB: 7PG9
PDB: 7V5G
PDB: 7V5M

**Figure S10 VCP and 20S cooperativity explained by the VCP C-terminal HbYX motif**

(**A, B**) Purified enzymes used in degradation assays. (**A**) Coomassie-stained gel of human 20S and 19S proteasomes. (**B**) Coomassie-stained gel of the human VCP, NPLOC4, and UFD1, along with ATPase activity assay in which NPLOC4/UFD1 (UN) without VCP served as background control (error bars: s.d. n > 3).
**(C)** Workflow of the cryo-EM SPA on the human 20S co-complexed with the VCP C-terminal peptide (DNDDDLYG). A representative micrograph, 2D class averages are shown. The 3D reconstruction is further classified without alignment and with a mask around the alpha subunits.
(**D-G**) Reconstructions from the cryo-EM SPA. (**D**) 35% of 20S contain no bound peptide, docked with PDB 7PG9 in ribbon representation. (**E**) 25% of 20S exhibit multiple signals in the inter-subunit pockets of PSMA (boxed) at a lower resolution; the reconstruction is docked with PDB 7PG9. (**F**) 16% of 20S docked with its own ribbon model with enlarged views of the boxed region in **(G)**, where the bound peptide is modeled as poly-alanine. The corresponding FSC curves, Euler angle distributions, and the local resolution maps of the top views including the inter-subunit PSMA pockets are shown for each class.
(**H**) AlphaFold3 prediction shows that all inter-subunit pockets between the human PSMA (cyan) can accommodate the VCP C-terminal peptide DNDDDLYG (green), resulting in a minor shift (RMSD: 0.49 Å) compared to the free human 20S (PDB 7PG9, magenta). The top view is displayed.
**(I)** Top view of the PDB models of the human free 20S (7PG9 blue) overlaid with processing 20S (7V5M green, 7V5G magenta).
Scale bar: 100 nm (**C**), 1 nm (**D-F**).
